# Emergence of the erythroid lineage from multipotent hematopoiesis

**DOI:** 10.1101/261941

**Authors:** Betsabeh Khoramian Tusi, Samuel L. Wolock, Caleb Weinreb, Yung Hwang, Daniel Hidalgo, Rapolas Zilionis, Ari Waisman, Jun Huh, Allon M. Klein, Merav Socolovsky

## Abstract

Red cell formation begins with the hematopoietic stem cell, but the manner by which it gives rise to erythroid progenitors, and their subsequent developmental path, remain unclear. Here we combined single-cell transcriptomics of murine hematopoietic tissues with fate potential assays to infer a continuous yet hierarchical structure for the hematopoietic network. We define the erythroid differentiation trajectory as it emerges from multipotency and diverges from 6 other blood lineages. With the aid of a new flow-cytometric sorting strategy, we validated predicted cell fate potentials at the single cell level, revealing a coupling between erythroid and basophil/mast cell fates. We uncovered novel growth factor receptor regulators of the erythroid trajectory, including the proinflammatory IL-17RA, found to be a strong erythroid stimulator; and identified a global hematopoietic response to stress erythropoiesis. We further identified transcriptional and high-purity FACS gates for the complete isolation of all classically-defined erythroid burst-forming (BFU-e) and colony-forming progenitors (CFU-e), finding that they express a dedicated transcriptional program, distinct from that of terminally-differentiating erythroblasts. Intriguingly, profound remodeling of the cell cycle is intimately entwined with CFU-e developmental progression and with a sharp transcriptional switch that extinguishes the CFU-e stage and activates terminal differentiation. Underlying these results, our work showcases the utility of theoretic approaches linking transcriptomic data to predictive fate models, providing key insights into lineage development *in vivo.*

## Introduction

Red cells are produced at a higher rate than any other blood lineage, increasing further during fetal development, hypoxia, bleeding and anemia. The abundance of erythroid progenitors in hematopoietic tissue provides a unique opportunity for dissecting how a multipotent progenitor differentiates into a single lineage in situ, a process of fundamental biological interest and of clinical relevance.

Erythropoiesis has two principal phases. During the erythroid terminal differentiation (ETD) phase^1–5^, a Gata1-driven transcriptional program remodels erythroid precursors into red cells through several well-described stages^6,7^. ETD is preceded by a much less well delineated phase of early erythropoiesis. Beginning with the hematopoietic stem cell (HSC)^8^, a series of poorly defined intermediates give rise to broad functional categories of unipotential erythroid progenitors, identified four decades ago by their formation of erythroid ‘bursts’ (burst-forming unit-erythroid, BFU-e) or smaller erythroid colonies (colony-forming unit-erythroid, CFU-e)^9,10^ in semi-solid medium. Although there has been some success in prospectively enriching for erythroid progenitors^11–16^, a direct, complete and high-purity isolation and molecular characterization of adult murine BFU-e and CFU-e progenitors from hematopoietic tissue has not been attained. More broadly, there have been no strategies that systematically identify the entire cellular and molecular trajectory of the early erythroid lineage as it first arises from the HSC and progresses to the point where the ETD program is activated.

Probing the earliest stages of erythropoiesis requires exploring how HSCs and multipotential progenitors (MPP) diversify into progenitors for each of the hematopoietic cell fates. Single cell approaches have recently upended established models of hematopoiesis, showing that populations of early progenitors that were thought to be similar in their developmental stage and fate potentials are in fact highly heterogeneous in both respects^17–25^. A number of alternative models have now been proposed, replacing the classic hematopoietic tree ^26–28^ with a ‘flatter’ hierarchy, in which uni-lineage or oligo-lineage progenitors derive directly from a heterogeneous set of lineage-biased HSCs or MPPs ^18–20,22,24^. The new modeling of hematopoiesis focuses particularly on cell transcriptional state, rather than cell fate, and is highly dependent on the tools employed in the analysis of high-dimensional cell states; these tools are currently undergoing intense innovation ^29–37^. Attempts to describe the structure of hematopoiesis have so far relied on clustering^19^, which may fail to capture continuum behaviors if they exist; diffusion maps ^37,38^, which are powerful for branching models but provide less detail of highly complex processes; and the ordering of progenitor trajectories based on their similarity to differentiated cell types^24^, which may overlook progenitors that do not resemble mature cells. Accurate reconstruction of developmental trajectories might also be compromised by biases introduced through the use of fluorescence-activated cell sorting (FACS), with restrictive FACS gates that exclude unknown cell types ^23^, and the loss of sensitive cell subsets during high-speed sorting ^39^. It is still not clear how to reconcile cell fate assays with cell state maps proposed from single cell profiling, and indeed this remains a general challenge in stem cell biology.

Here we investigated the derivation of erythroid progenitors from multipotential progenitors, and their subsequent development *in vivo.* We undertook single-cell RNA-seq (scRNA-seq) of a broad set of freshly isolated hematopoietic progenitors selected only through their expression of a single cell surface marker, Kit, using the InDrops platform^40^. We applied an analytical tool, Population Balance Analysis (PBA), that predicts fate probabilities from static snapshots of single cell transcriptomes through dynamic inference, which allowed us to define a FACS strategy to isolate cells in progressive stages of early erythropoiesis. Using single cell fate assays, we then confirmed a number of detailed predictions regarding the early hematopoietic hierarchy and erythroid developmental progression. The insights obtained regarding early erythropoietic fate control may be applicable to other differentiation models, and include novel erythropoietic regulators with potential translational relevance.

## Results

### Single-cell RNA-seq of Kit+ hematopoietic progenitors

To investigate early erythropoiesis from multipotency to ETD, we performed scRNA-Seq on hematopoietic progenitor cells (HPCs) isolated from murine adult (8 weeks) bone marrow (BM). HPCs were isolated using magnetic beads, conditional on expression of the cell-surface receptor Kit, expressed on all hematopoietic stem and early progenitor cells^41,42,43^ (**Fig. 1a**). This inclusive approach preserves the relative abundance of the various early progenitor cell states, allowing a largely unbiased reconstruction of early hematopoietic differentiation. After filtering scRNA-Seq data, we carried forward 4,763 HPC transcriptomes for analysis (see **Data Availability** for interactive tools).

**Figure 1.**
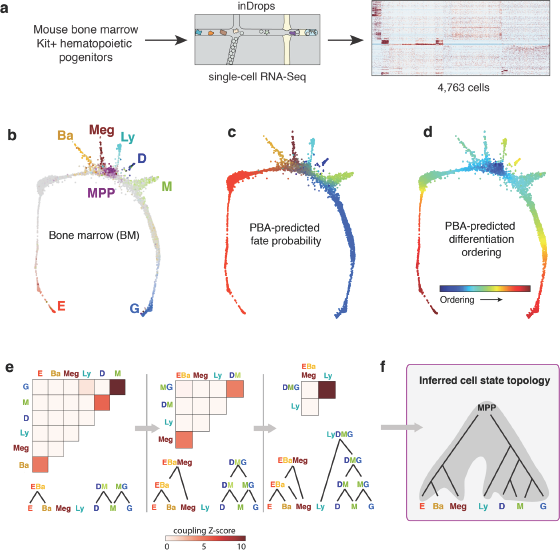
Prediction of the early hematopoietic hierarchy by single-cell RNA-seq. **a** Schematic of the experiment profiling Kit+ mouse bone marrow cells using scRNA-seq. **b** Visualization of single-cell transcriptomes using a SPRING plot (each point is one cell). Colors indicate composite expression level of lineage-specific genes. E, erythroid; Ba, basophilic; Meg, megakaryocytic; Ly, lymphocytic; D, dendritic cells; M, monocytic; G, granulocytic; MPP, Multipotential progenitors. **c,d** Parameterization of the cell state graph using Population Balance Analysis (PBA), an algorithm that describes the graph position of each cell by a set of computationally-predicted fate probabilities (c) and a pseudo-temporal ordering with MPPs at the origin and terminating with the most mature observed cells of each lineage (d). Each lineage probability (E, Ba, Meg, Ly, D, M, G) is assigned a color as in (b). **e,f** A cell state hierarchy encodes the cell graph topology. Lineage-biased states were identified by calculating the fraction of PBA-predicted cells with bilineage coupling for every two lineages, and comparing to what would be expected after fate randomization (e). Iteratively joining fates based on their pairwise coupling produced a final hematopoietic multipotent cell state hierarchy (f).

We used the SPRING algorithm^44^ to visualize the single-cell transcriptomes (**Fig. 1b**). SPRING represents scRNA-Seq data as a graph of cells (graph nodes) connected to their nearest neighbors in gene expression space, projected into two dimensions using a force-directed graph layout. This unsupervised visualization suggests that HPCs occupy a continuum of transcriptional states, rather than discrete metastable states, a result which contrasts with single cell data from mature blood lineages and is supported by formal tests of graph interconnectivity (**Extended Data Fig. 1a**). The nature of the HPC continuum becomes apparent when SPRING plots are colored based on expression of lineage-specific markers (**Fig. 1b, Extended Data Table 1**) the cells are found to organize around an undifferentiated core, from which seven distinct branches emerge corresponding to progenitors of the granulocytic (G), monocytic (M), dendritic (D), lymphoid (Ly), megakaryocytic (Meg), basophilic/Mast cell (Ba), and erythroid (E) lineages. Observing this structure depends critically on collecting cells using a broad selection marker. Previous studies^19,23^ which instead combined cells from multiple classically-defined sub-populations, distort the relative abundance of progenitor cell states but are revealed by SPRING to encode the same lineage relationships (**Extended Data Fig. 1b**).

### The transcriptional state continuum of HPCs is hierarchical, but not a strict tree

Early work proposed hematopoietic hierarchies in which MPPs first give rise to oligopotential progenitors, which in turn differentiate into unipotent progenitors^26–28^. In contrast, recent single cell approaches have suggested a flat hierarchy with cells transitioning directly from MPPs to unipotent progenitors^18–20,22,24^. To understand how these models might relate to our observed cell state topology, we developed an approach for studying single cell continua, called Population Balance Analysis (PBA)^45^ which maps each cell to a low-dimensional space that encodes the cell graph topology in the form of predicted cell fate probabilities. For each hematopoietic progenitor, PBA defines seven putative commitment probabilities that encode graph distances to each of the observed terminal fates (**Fig. 1c, Extended Data Fig. 1c**), as well as the distance from the undifferentiated CD34^hi^/Sca1^hi^ MPPs (**Fig. 1d**). To determine whether the PBA map predicts the existence of bipotential and oligopotential progenitors in the hematopoietic hierarchy, we computed a coupling score that determines whether any two fate potentials are co-expressed in the same progenitor, at rates higher than expected by chance (**Fig. 1e**). A transcriptional state hierarchy was then formalized by identifying correlated pairs of terminal fates, iteratively joining fates until a multipotent state is reached (**Fig. 1e,f**).

The topology revealed by this analysis firmly supports the hierarchical view of hematopoiesis. Specifically, it shows that MPPs diverge into progenitors with correlated E/Meg/Ba fates, or correlated Ly/Myeloid fates (**Fig. 1e**). The existence of correlated sets of terminal fates argues against the notion of a flat state hierarchy. However, the transcriptional-state hierarchy emerges from correlations on a continuum, rather than from discrete oligopotent progenitors, and it predicts some refinements over current models, particularly two key features: first, the erythroid fate is correlated with the basophil/mast cell fates, in addition to the known correlation with the megakaryocytic fate. Second, among myeloid progenitors we identify dendritic-monocyte (DM) and granulocytic-monocyte (GM) coupling, but no corresponding DG coupling, indicating that monocyte differentiation may occur through two distinct molecular trajectories and not through a strict tree-like branching process.

By formalizing the HPC transcriptional hierarchy, PBA allows examination of gene expression correlating with putative cell fate probability. Here we focus on erythroid differentiation, but we provide a global summary of the transcription factors, receptors and chromatin modifiers correlating with each fate choice (**Extended Data Fig. 2, Extended Data Table 2**).

### scRNA-Seq-guided isolation of putative erythroid progenitors

To test the computational predictions of fate couplings (**Fig. 1e-f**), and to define early erythroid progenitors, we developed a flow cytometric sorting strategy to isolate the hematopoietic subpopulations defined by scRNA-Seq. Guided by the single cell expression patterns, we divided Kit+ bone-marrow cells based on cell surface CD55, which was recently shown to mark progenitors biased for the Meg/E fates^17^. We then subdivided Kit^+^CD55^+^ cells into P1 to P5, by combining the alpha integrin CD49f *(Itga6)* with markers previously used to identify Meg/E-biased progenitors ^11,17,15^ (**Fig. 2a, Extended Data Fig. 3**). Using qRT-PCR for genes that mark specific locations within the SPRING graph (**Extended Data Fig. 4**), and scRNA-Seq analysis (11,241 cells post-filter) (**Fig. 2b**), we mapped cells from each of the sorted subpopulations back to regions of the SPRING graph. We found that P1 and P2 represent high-purity sub-populations on the putative erythroid branch, with P1 predicted to be committed, and P2 mostly committed, to the erythroid fate (**Fig. 1c**); P3 and P4 are enriched for the basophilic and megakaryocytic branches, respectively, and P5 contains oligopotent and multipotent cells with an erythroid fate bias. Some MPPs, myeloid and lymphoid states were within the CD55^-^ zone of the plot. Interestingly, with a relatively large number of cells sampled from small parts of the Kit+ continuum, the P3 subpopulations bifurcated into branches with markers indicative of either the basophil or mast cell lineages (**Extended Data Fig. 5**).

**Figure 2.**
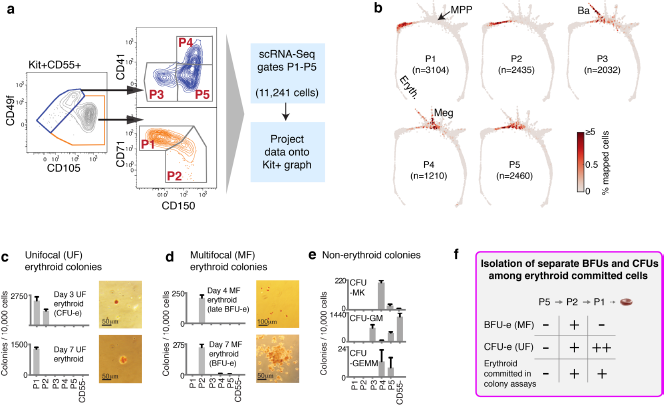
A novel sorting scheme isolates high purity progenitors of early erythropoiesis. **a** Flow-cytometric sorting strategy, dividing Kit+CD55+ bM cells into gates P1 to P5 based on expression of CD49f *(Itga6),* CD105 (Eng), CD41 *(Itga2b),* CD150 *(Slamf1),* and CD71 *(Tfrc).* Freshly sorted cells from each gate were profiled using scRNA-Seq. **b** The sorted subpopulations P1-P5 are localized on the Kit+ graph by assigning single cell transcriptomes from each sub-population to their most similar counterparts. The density of projected cells is represented using a heatmap. **c-e** Colony formation assays in methylcellulose carried out on freshly sorted P1 to P5 subsets and on Kit+CD55^-^ cells show that P1 and P2 subsets completely define classic CFU-e and BFU-e progenitors. Erythroid colonies were classified as either unifocal (c) or multifocal (d). CFU-MK=Megakaryocytic colonies (see **Extended Data Fig. 6**), CFU-GM= granulocytic and/or monocytic colonies; CFU-GEMM= mixed myeloid colonies. Results are mean±SE of two independent sorts. Colony assays were performed in triplicate. Images show erythroid colonies, stained with diaminobenzidine to highlight hemoglobin expression. The P1 population enriches 150-fold for CFU-e, compared with total BM; the P2 population enriches 50-fold, 80fold and 45-fold for CFU-e, late and early BFU-e, respectively. **f** Summary of erythroid colony formation potential in FACS subsets.

### Fate assays define early erythroid, basophilic and megakaryocytic progenitors

We examined the differentiation potential of each of the sorted populations P1-P5, and by extension, the predicted fate probabilities for defined transcriptional states (**Fig. 1c**). Using methylcellulose colony forming assays, no subset other than P1 and P2 gave rise to erythroid colonies (**Fig. 2c-d**), and conversely, P1 and P2 gave rise to no other colony type scored [CFU-GM^46^, mixed Myeloid / Erythroid /Megakaryocytic (CFU-GEMM^47–49^), and megakaryocyte colonies (CFU-Mk^50^)] (**Fig. 2e**), suggesting that, in this assay, P1 and P2 contain all of the unipotential erythroid progenitors of the bone-marrow and few other progenitor types. Of interest, P1 colonies were small (30-100 cells) and unifocal (**Fig. 2c**), maturing on day 3 (CFU-e) or later, whereas P2 colonies were largely multifocal, maturing on day 4 (late BFU-e) or later (early BFU-e) (**Fig. 2d**). Thus, we confirmed that P1 cells were indeed closer to erythroid maturation than P2 cells, consistent with their location on the cell state graph (**Fig. 2b**); and we found that the molecular stage of progenitors determines their ability to form either multifocal or unifocal colonies.

In contrast to P1 and P2, the less differentiated P5 subset gave rise to CFU-GM, CFU-GEMM and some CFU-Mk, in agreement with its location in the cell state graph. Also as expected, P4 was enriched for megakaryocytic colonies (**Fig. 2e, Extended Data Fig. 6a**).

To obtain a more refined view of HPC fate potential, we sorted single cells from each of the Kit+ progenitor sub-populations into liquid culture wells in the presence of a cocktail of cytokines that support differentiation of myeloid and erythroid lineages (**Fig. 3a**), and assayed their clonal output after 3 (for P1), 7 or 10 days by multi-parameter FACS, using markers of erythroid (“E”, Ter119+), basophil (“Ba”, FceR1a+), megakaryocytic (“Meg”, CD41+), granulocytic (“G”, Gr-1+) and monocytic (“M”, Mac-1+) lineages. We quantified these five lineages in 1,158 single cell clones (**Fig. 3b, Extended Data Fig. 6b**). We found that unipotential clones for the E, Ba, Meg and G/M lineages largely originated in the P1/P2, P3, P4 and CD55-subpopulations, respectively, consistent with their location on the SPRING plot (**Figs 3b, 1c, 2b**). In addition, many of the Kit+ clones contained multiple lineages; we found strong, statistically significant couplings between the E, Ba and Meg cell fates on the one hand, and the G and M fates on the other (**Fig. 3c**; z score absolute value > 10 compared to randomized data), consistent with both known (E/Meg, G/M) and novel (E/B) cell fate correlations predicted by PBA analysis (**Fig. 1c,e-f**). In addition, we expected that progenitor clones containing both E and Ba cells would be depleted in CD55-cells and enriched in sub-populations P2-P5, which map close to the E/B branch point in the scRNA-Seq data (**Figs. 2b, 1c**), and indeed this was confirmed by the single cell clonal assays (**Fig. 3d**). The same results are reflected in bulk liquid culture assays (**Extended Data Fig. 6c**), which show that the P2 fraction largely gives rise to erythroid cells, but also to Meg (CD41+) and Ba (FceR1a+) progeny (**Extended Data Fig. 4a-b**). We conclude that the E/B/Meg fates are coupled both transcriptionally and functionally, while being anti-coupled to the G/M fates, and that scRNA-Seq data can be used to generate successful predictions of HPC states and fates.

**Figure 3.**
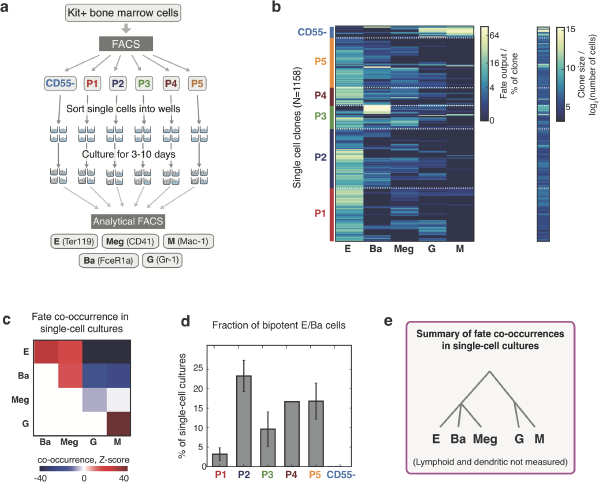
Evidence of predicted fate couplings through single cell *in vitro* fate assays. **a** Schematic illustration of liquid culture assay for measuring clonal lineage output. E, erythroid; Ba, basophil; Meg, megakaryocyte; G, granulocyte; M, monocyte. **b** *Left:* Heat map of clonal lineage output. Rows represent single clones, with colors indicating the fraction of cells belonging to each measured lineage. *Right:* Heat map of the total number of cells produced by each clone. **c** Heat map of lineage co-occurrences observed in the single cell fate assays. Co-occurrence was computed by comparing the number of clones producing a pair of fates (≥2% of cells in a clone) to the number expected after randomizing fates. **d** Average proportion of bipotent erythroid-basophil (EBa) clones from each FACS gate establish that P2 sits near the EBa branch point. Clones were considered bipotent if E and Ba cells, but no other fates, each comprised >2% of the total number of cells. Error bars are SE of the unweighted mean of two (P1, P2, P3) or three (P5) independent sorts. Only a single sort was performed for P4 and CD55-. **e** Cell state hierarchy based on the fate co-occurrence in single-cell cultures.

### The erythroid differentiation trajectory

Based on the structure of the scRNA-Seq data and the fate potential assays, we partitioned the continuum of cell states between MPPs and ETD into three stages (**Fig. 4a**). We termed these: (1) erythroid-basophil-megakaryocytic progenitors (EBMegP), (2) early erythroid progenitors (EEP), and (3) committed erythroid progenitors (CEP). EBMegPs are oligopotent cells near branch points to megakaryocytic and basophil lineages, that are biased away from the G/M fates. They are strongly represented in the P5 and P2 subpopulations (**Fig. 2b**). The EEP stage is a narrow region of the graph, just past the final non-erythroid fate branch point. It forms most of the P2 gate, and functionally corresponds to BFU-e progenitors (**Fig. 2b-d**). The CEP stage contains the majority of pre-ETD unipotential erythroid progenitors, and functionally corresponds to CFU-e progenitors (**Fig. 2b-d**). These stage definitions integrate the results of **Figs. 2–3** to reconcile fate, FACS gates and scRNA-Seq-based definitions of differentiation progression.

**Figure 4.**
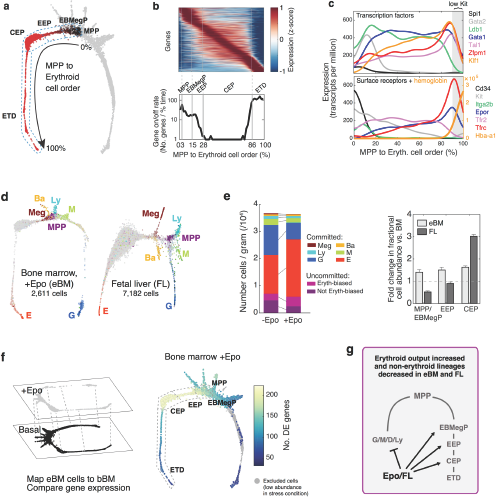
Stages of early erythropoiesis and the global erythroid stress response. **a** Definition of three transitional stages between multipotency (MPP) and erythroid terminal differentiation (ETD), based on scRNA-Seq and fate assays of FACS subsets P1-P5. EBMegP: Erythroid-Basophil-Megakaryocyte biased progenitors; EEP: Early Erythroid Progenitors, encompassing all BFU-e and some CFU-e; CEP: Committed Erythroid Progenitors, which are pure CFU-e. The SPRING plot shows the erythroid fate probability predicted computationally by PBA. **b** *Top:* Heat map of dynamically varying genes (rows) in cells (columns) arranged by the graph ordering of cells from MPP to ETD. Gene expression smoothed using a Gaussian kernel and plotted in order of peak expression position. *Bottom:* Density of expression inflection points (number of genes turning on or off), indicating the rate of change of gene expression with progression from MPP to ETD. The x-axis represents the PBA-predicted differentiation ordering of cell transcriptomes, from the least differentiated cell at 0% to the most mature cell at 100%, with remaining cells uniformly spaced in between. **c** Gene expression traces for established erythroid regulators along the ordered erythroid trajectory (region with low Kit expression, highlighted in grey). **d** SPRING plots of scRNA-Seq data from erythroid stress conditions show a strong conservation of Erythroid-Basophil-Megakaryocyte coupling, with major expansion of the erythroid lineage. Colors show composite cell type specific gene expression as in **Fig. 1b**. **e** *Left:* Absolute quantification of the number of Kit+ cells in the mouse (**Extended Data Fig. 10**) reveals that the expansion of the erythroid lineage after Epo stimulation occurs at the expense of all other lineages, except for the Basophilic lineage. Among uncommitted cells, the number of predicted erythroid-biased progenitors (cells included in the MPP to Erythroid ordering) increased while the remainder diminished. Committed cells were defined by a PBA-predicted erythroid fate probability >0.5. *Right:* FL erythropoiesis expands primarily in CEP, while Epo leads to expansion of even the earliest erythroid progenitor cell states. **f** Epo response leads to gene expression changes all along the early erythroid trajectory, seen by mapping cells from the stress sample (eBM) onto the basal BM and then conducting a differential gene expression analysis. The heatmap shows the number of genes differentially expressed (DE) between mapped stress and basal cells. **g** Summary of changes in cell state abundance during stress erythropoiesis.

To establish the transcriptional events of the erythroid differentiation trajectory, we computationally isolated cells lying on the PBA-predicted state progression from MPP to ETD, creating a smoothed time series for every gene, akin to previously published pseudotemporal ordering algorithms^51,52,53^ (**Fig. 4b**). The expression of known erythroid regulators recapitulated their expected dynamics (**Fig. 4c**): transcription factors *Gata2*^54^, Tall^55^, Ldb1^56^ or *Spi1* (PU.1)^57^, were expressed at the MPP starting point; the erythroid transcription factor *Gata1* was induced early, concurrent with suppression of *Spi1* (PU.1) and Gata2^58^. The Epo receptor, *EpoR*^59^, was also induced early, rising to a further peak at the transition to ETD, identifiable by the sharp induction of erythroid genes such as a-globin *(Hba-a1).* We validated expression of canonical TFs in the sorted populations P1-P5, including the early expression of *Gata1* mRNA and protein in EEP/CEP (**Extended Data Fig. 7a,b**). We further established that a graded increase in *Tfrc* (CD71) expression is a reliable marker of continuous progression through the EEP and CEP stages, by showing that transcriptomes of sorted CD71^high^ P1 cells map to late CEP stage (N=752 cells post-filter, **Extended Data Fig. 7c**), and by finding that CD71 expression gradually increases in sorted P2 and P1 cells differentiating *in vitro* (**Extended Data Fig. 7d**). A further, sharp increase in CD71/Tfrc expression takes place at the transition from CEP to ETD (**Fig. 4c**)

In total, >4,500 genes varied significantly along the erythroid trajectory (**Extended Data Table 3**). Of these, the most striking dynamic was by a large group of genes that were induced at the onset of the CEP stage, and were suppressed during a second, sharp transcriptional switch at the CEP/ETD transition (**Fig. 4b**). By performing a gene set enrichment analysis (GSEA) on dynamic gene clusters (**Extended Data Figs. 7e,8 and Extended Data Table 4**), we found that the most dominant clusters were of genes specifically upregulated at the CEP stage; these were enriched for cell cycle and growth-related genes, including mTOR signaling, nucleotide metabolism, RNA processing, oxidative phosphorylation, DNA replication, and mitochondrial components. These pathways suggest that one of the functions of the CEP stage is to act as an “amplification” module, a hypothesis supported by the finding that CEPs are the most abundant cells in early erythropoiesis. GSEA also predicted several new epigenetic and transcriptional regulators of erythroid progression (**Extended Data Fig. 9 and Extended Data Table 4**), and interestingly, showed that while Gata1 is expressed from the start of the CEP stage, the vast majority of its canonical erythroid targets are induced only at the transition from CEP to the ETD stage. Taken together, the temporal ordering of the single-cell transcriptomes accurately recapitulates known events of early erythropoiesis and uncovers a dedicated CEP transcriptional program that is distinct from the ETD program.

### Stress generates erythroid-trajectory-wide changes but preserves the structure of the hematopoietic hierarchy

We asked whether erythropoiesis occurring in different contexts from the homeostatic bone marrow would still show the same gene expression branching topology, as well as transitional EEP and CEP states. Fetal liver (FL) and Epo-stimulated red cell production in the bone marrow (eBM) are both examples of accelerated, or stress, erythropoiesis. In mid-gestation FL, erythropoiesis is rate limiting to fetal growth, accounting for >95% of the tissue^60^. In the BM, Epo increases erythropoietic output by promoting the survival of CFU-e/ proerythroblasts^61,62^. Whether Epo exerts effects upstream of the CEP/CFU-e stage is unknown.

We therefore performed unsupervised scRNA-Seq analysis in FL from mid-gestation (E13.5) embryos (*N*=7,182 cells post-filter), and in eBM (*N*=2,611 cells post-filter), isolated following 48 hours of Epo stimulation *in vivo,* and compared the results to baseline BM data. The scRNA-Seq data, shown through two unsupervised SPRING graphs in **Fig. 4d**, revealed a remarkable conservation of the key features of the hematopoietic hierarchy and erythroid differentiation in particular. In both cases, basophils and megakaryocyte branches emerged just prior to erythroid, with reduced lymphoid and myeloid branches diverging even earlier.

In both stress conditions and particularly in FL, the overall proportion of cells in the erythroid trajectory increased compared with basal BM (**Figs 4d,e**). In FL, the increase was predominantly at the CEP stage, whereas, surprisingly, in eBM all pre-ETD cells in the erythroid trajectory increased in abundance, including uncommitted MPPs and EBMegPs. The absolute number of Kit+ cells in eBM did not change (**Fig. 4e** and **Extended Data Fig. 10**), suggesting that the increase in abundance of erythroid trajectory cells occurs at the expense of other lineages. A number of mechanisms could account for such an effect, including altered intrinsic fate bias of MPPs ^63,64^, extracellular-signal-mediated amplification of MPPs and EBMegPs with pre-existing erythroid bias, and/or inhibitory influences on downstream non-erythroid progenitors.

Analysis of Epo-induced differentially-expressed genes showed the largest number in EEP and CEP, but also showed changes in gene expression in EBMegP and MPPs (**Fig. 4f**). Notably, these included downregulation of targets of C/EBPβ, a TF that biases differentiation away from erythroid/megakaryocytic fates and towards granulocytes/monocytes^65^. We also saw a large number of differentially-expressed genes in the FL (**Extended Data Fig. 11, Extended Data Table 5**). In both FL and eBM, differentially-expressed genes included new, as well as known, regulatory changes, such as the upregulation of Stat5 targets *(Cish, Socs3, Trib3)*^3,66,67^, *Podxl*^68^, ROS pathway genes^3,66,69^ and PU.1 targets^32^. For each of the known or novel stress responses, our analysis predicts its precise localization within the erythroid trajectory.

Taken together, our observations show that the cell state branching structure is maintained during accelerated erythropoiesis. We identified extensive changes in gene expression and in cell abundance in response to Epo, in MPPs and throughout the ensuing erythroid progression, well beyond the currently known mechanism of Epo-driven erythropoietic expansion^61,62^.

### Novel growth factor regulators of early erythropoiesis

In the scRNA-Seq data, we screened EEP and CEP genes that encode cell-surface receptors with known ligands, which could be tested for a potential regulatory effect. We identified three such receptors: *Ryk, Mst1r* and *Il17ra* (**Fig. 5a, Extended Data 12**). Ryk functions in non-canonical Wnt signal transduction^70^. It was cloned in bone-marrow CFU-e, but its function there remained unknown^71^. Mst1r (macrophage stimulating 1 receptor/ RON kinase) is a receptor-tyrosine kinase (RTK) expressed in multiple tissues and tumors^72^. It was found to associate with the EpoR in a CFU-e culture system, but could not be activated directly by its ligand, Macrophage Stimulating Protein (MSP/Mst1), in that context^73^. *Il17ra* encodes one of several receptors for the IL-17 proinflammatory cytokines^74^, found to be broad inhibitors of bone-marrow progenitors^75,76^. However, the expression of an IL-17 receptor by erythroid progenitors had not been documented.

In BM, *Il17ra* was expressed by EEPs/P2 and to a lesser extent by CEPs/P1, in addition to non-erythroid cells. Both *Ryk* and *Mst1r* are expressed more selectively, though not exclusively, by EEP and CEP cells (**Fig. 5a, Extended Data Fig. 12a-b**). We stimulated Ryk, Mst1r and IL-17Ra with their respective ligands, Wnt5a, MSP and IL-17a, using Epo-dependent erythroid colony formation as readout (**Fig. 5b,c, Extended Data Fig. 12c**). Epo concentration in blood varies from ~10 mU/ml in health, to 10,000 mU/ml in severe anemia^77^. We found that addition of MSP to FL in the presence of mildly elevated Epo (50 mU/ml) was equivalent to a 10-fold increase in Epo concentration, generating over 2-fold the number of CFU-e colonies. However, MSP was inhibitory in other contexts, and Wnt5a was a consistent and potent inhibitor of all erythroid colony formation in both FL and BM (**Fig. 5b, Extended Data Fig. 12c**). By contrast, IL-17a mediated a striking potentiation of adult BM CFU-e colony formation throughout the Epo dose-response curve, giving rise to 4-fold more colonies at lower Epo (50mU /ml), and to a ~50% increase in colony formation when added to maximal levels of Epo (**Fig 5b,c**). It had a milder stimulatory effect in BFU-e progenitors, and no effect in FL (**Extended Data Fig. 12**).

**Figure 5.**
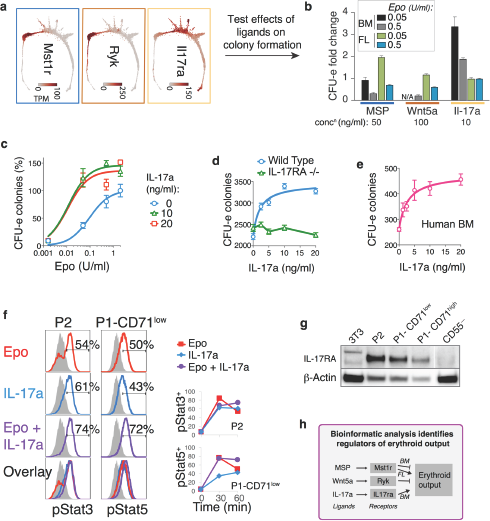
Identification and validation of growth factors regulating early erythropoiesis. **a** Expression patterns for *Mst1r, Ryk* and *Il17ra* show peaks at CEP or EEP (See also **Extended Data Fig. 12a**). qPCR validations on FACS-sorted population are provided in **Extended Data Fig. 12b**. **b** Representative changes in the number of CFU-e colonies formed in methylcellulose in the presence of adding the cognate ligands MSP, Wnt5a or Il17a to culture media. Error bars show SD of two independent experiments, with four replicates per experiment. **c** Representative Epo dose response curve for IL-17a, showing a persistent IL-17a response even at saturating doses of Epo. Dose response curves in BM and FL for all ligands are provided in **Extended Data Fig. 12c**. **d** IL-17a response is dependent on IL-17Ra expression, seen by loss of response in colonies from IL-17Ra-/-mice. Colonies scored per 500,000 plated BM cells in the presence of Epo at 0.05 U /ml. Data is mean ± SD of triplicate samples; representative of two independent experiments. **e** IL-17a also stimulates CFU-e in freshly isolated human bone marrow mononuclear cells. Colonies scored per 85,000 cells plated. Data is mean ± SD of triplicate samples. **f** FACS analysis of fixed and permeabilized cells stained for pStat3 and pStat5 shows that IL-17a rapidly activates intracellular signaling through these pathways independently of Epo. Freshly isolated mouse BM cells were starved of cytokines for 3 hours, and then stimulated with either Epo, IL-17a or both cytokines. FACS profiles on *left* are for baseline (starved) cells (in grey), and 60 minutes-post stimulation (in color). *Right:* Time course of the fraction (%) of cells positive for either pStat in the same experiment. Representative of two experiments. **g** Western blot show IL-17Ra peak expression in P2/EEP cells dropping in P1/CEP and in the granulocytic branch (which contributes most of the CD55-cells), consistent with the SPRING plots in (a). h Summary of effects of novel growth factor regulators on erythroid output.

The stimulatory effect of IL-17a on CFU-e formation was specifically mediated through expression of IL-17Ra, as shown by its absence in BM of IL-17Ra-deleted mice (**Fig 5d**), and was also evident in human BM (**Fig 5e**). Further, IL-17a stimulation was saturable, with a low EC_50_ [2.0 ± 1.2 ng/ml (60 pM) in mouse, 2.7 ±1.6 ng/ml (81 pM) in human BM], consistent with it being the direct consequence of high affinity-binding of IL-17a to IL-17Ra. Stimulation with IL-17a resulted in rapid phosphorylation of the intracellular signaling mediators Stat3 and Stat5 in CEP and EEP (**Fig. 5f**). Further, freshly sorted BM CEP/P1 and EEP/P2 expressed IL-17Ra by western blotting (**Fig. 5g**). Taken together, our findings suggest previously unknown complex regulatory modulation of EEP and particularly of CEP through the expression of a number of growth factor receptors new to erythropoiesis.

### Extensive remodeling of the cell cycle during erythroid developmental progression

In a final analysis, we asked what governs progression through the CEP stage and its termination at the transition to ETD. We previously reported that ETD activation in the FL is a cell-cycle synchronized event that takes place within the span of a single S phase, and is dependent on S phase progression ^60^; further, this S phase is unique in that it is shorter and faster than S phase in pre-ETD cells^78,79^. These conclusions, based on experimental analysis of large FL sub-populations, predict that exit from CEP should show a clear S phase signature. In our scRNA-Seq data, we found that during CEP/ETD transition, genes whose expression marks specific cell cycle phases indeed form an orderly sequence of close, sharp peaks, corresponding to G1/S, S, G2 and G2/M, likely representing a single cell cycle (**Fig. 6a-b**). This and the following results hold even when cell cycle genes are omitted for ordering the erythroid trajectory (**Extended Data Fig. 13a-c**). Significantly, by reversibly inhibiting DNA replication in S phase using the DNA polymerase inhibitor aphidicolin^80^, we found that the CEP/ETD switch in adult BM was not only synchronized with, but also dependent on, S phase progression (**Extended Data Fig. 13d-f**).

**Figure 6.**
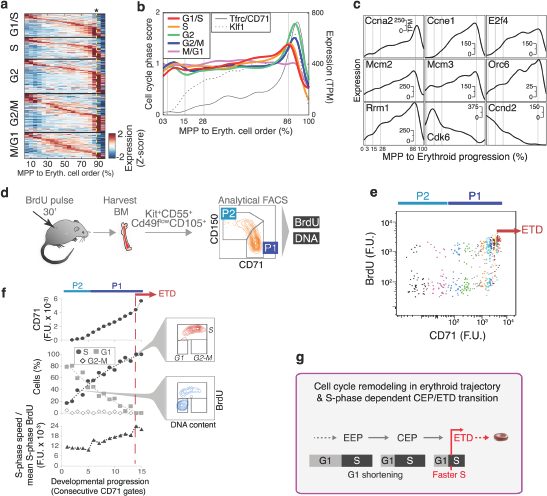
Erythroid progression correlates with progressive remodeling of the cell cycle, culminating with an S phase-dependent activation of the erythroid terminal differentiation program. **a** Heatmap of cell cycle-phase specific genes^88^, ordered by their peak expression along the MPP to erythroid progression, reveals synchronization of cell cycle at the CEP/ETD transition (indicated by *). **b** T races of the mean expression of all the genes specific to each cell cycle phase along the MPP to erythroid progression (as in panel “a”). Activation of ETD takes place with sharp Tfrc/CD71 upregulation (black curve). See also **Extended Data Fig. 13h** for corresponding western blot analysis. **c** Representative cell cycle genes correlating or anti-correlating with progression through the CEP stage. **d** Schematic illustration of cell cycle analysis of erythroid progenitors *in vivo.* Mice (6-8 weeks) were injected with BrdU (100 μ of 10 mg/ml stock in PBS); BM was harvested at 30 min post injection, immediately fixed, and analyzed for BrdU incorporation and DNA content in P1 and P2 subsets, defined as in **Fig. 2a**. **e** Cell cycle distribution with developmental progression. BrdU incorporation in P1 & P2 cells is a function of CD71 expression. Cell coloring indicates their division into consecutive 7-percentile gates of increasing CD71 for use in panel “f”. See also **Extended Data Fig. 7c,d**, for evidence that CD71 levels mark developmental progression through the EEP/P1) and CEP/P2 stages. The transition to ETD is marked, at CD71 window where CD71 increases sharply, and all cells are synchronized in S phase (BrdU+) (see also f below) **f** CD71 expression (top), cell cycle phase distribution (middle), and intra-S phase DNA synthesis rate (lower panel), for all gates shown in panel “e”. Insets show representative FACS plots used to measure the cell cycle phase distribution. See **Extended Data Fig. 13g** for similar analysis in eBM and FL. **g** Summary of cell cycle changes during early erythropoiesis. Following a shortening of G1 phase during progression through the CEP stage, cells undergo a fast S-phase required for the transition to erythroid terminal differentiation.

The scRNA-seq data also revealed that changes to cell cycle machinery occur earlier and throughout CEP, perhaps in preparation for the switch to ETD. The genes whose expression most closely correlate with CEP progression (**Extended Data Table 6**) are enriched for Gene Ontology terms associated with cell cycle and DNA replication. Strikingly, regulators of S phase and the G1/S transition increase steadily through the CEP stage, and include *Cyclin E1 (Ccne1), Cyclin A2 (Ccna2), E2f4,* Mcm helicase subunits *(Mcm2-7), Orc6.* Conversely, the G1 phase regulators, *Cyclin D2 (Ccnd2)* and *Cdk6,* decrease steadily (**Fig. 6c**). To investigate these findings, we made use of the graded increase in CD71 as a marker of progression through the EEP/CEP stages (**Extended Data Fig. 7c-d**). We pulsed mice *in vivo* with the nucleotide analog BrdU in order to mark S phase cells, and then analyzed the cell cycle distribution within narrow consecutive windows of increasing CD71 expression in the P1 and P2 subpopulations (**Fig. 6d,e**). We found a graded but dramatic increase in the fraction of cells in S phase, while the number of G1 cells correspondingly decreased (**Fig. 6f**). The same results held in eBM and in FL (**Extended Data Fig. 13g**). There was no significant change in S phase speed/length, evidenced by intra-S phase BrdU signal ^78^ (**Fig. 6f**); suggesting that cells spend more time in S phase as a result of G1 shortening. A western blot of freshly sorted EEP/P2 and two developmentally-consecutive fractions of CEP/P1 further confirmed the increasing expression of key S phase regulators with developmental progression (**Extended Data Fig. 13h**). Taken together, our data suggest that progression through the erythroid trajectory is associated with extensive remodeling of the cell cycle phases (**Fig. 6g**).

## Discussion

Single-cell depictions of stem and progenitor cell hierarchies face a challenge in reconciling cell fate assays, molecular state maps, and functional responses to perturbation from homeostasis. Our scRNA-seq analysis reveals that HPCs occupy a continuum of transcriptional cell states, branching towards 7 distinct cell fates. Certain cell fate potentials are correlated, supporting a hierarchical view of hematopoiesis, with MPPs diverging either towards myeloid and lymphoid fates, or towards the erythroid/ megakaryocyte/ basophil cell fates. Yet unlike the classical models of hematopoiesis, HPCs do not separate into discrete and homogenous stages. The coupling of specific cell fates in early progenitors, which we validated with single-cell fate assays, is a critical feature by which our model differs from recent models of hematopoiesis, where uni-lineage progenitors arise directly from MPPs. It explains historical interpretations of the hematopoietic hierarchy that were based on fate assays of FACS-gated populations, which enrich for the fate couplings of a population’s constituent progenitors. Thus, our model’s predictions explains the appearance, at a population level, of the split of MPPs into E/Meg and LMPP branches ^28^, but further modify this by the prediction, validated experimentally, of a novel coupling between the basophil/mast cell and erythroid fates, recently also predicted by Velten et al. ^24^. This coupling is consistent with the expression of TFs GATA2 and Tal1 early in both lineages ^81,82^, and with recent reports that document the origins of mast cells and basophils as distinct from that of neutrophils ^83,84^. However, the continuum nature of the scRNA-Seq data does not rule out the existence of discrete epigenetic or signaling states among HPCs; such discrete states could give rise to a transcriptional continuum if their lifetime in single cells was comparable, or shorter than, the lifetime of mRNA molecules (hours to ~1 day).

We delineated the continuous differentiation trajectory of the erythroid lineage, from its origins in MPPs, through EBMegPs, with correlated erythroid, basophil/mast cell and megakaryocytic fate potentials, to unipotential erythroid progenitors. We identified the long-sought cytometric profiles and transcriptional states that correspond to the classical BFU-e and CFU-e erythroid progenitors in murine BM, which we named EEP and CEP, respectively. The dominant CEP stage has a dedicated transcriptional program, distinct from that of ETD, and is likely to function as a regulatory module controlling erythropoietic output, as evidenced by its expansion in stress, and by our identification of three novel growth factor receptors that regulate CEP/CFU-e numbers. These include strong stimulation by the pro-inflammatory IL-17Ra, possibly contributing to the growing complex interplay between erythropoietic output and inflammatory response pathways that underlie anemia of chronic disease ^85^. We further identified the cell cycle as a key process in both the progression, and termination, of the CEP stage. Cells shorten the G1 phase and spend increasingly more time in S phase as they progress through the CEP stage; the transition from CEP to ETD takes place as a rapid, S-phase dependent transcriptional switch, during the span of a single cell cycle in which S phase also becomes shorter. Of note, GATA-1 and other erythroid TFs that drive ETD are induced early in the EEP stage. We speculate that the extensive modulation of the cell cycle might provide additional context necessary for transcriptional activation of ETD, with likely implications for developmental progression of other lineages. The context provided by the cycle is unknown but may be related to recent findings of cell-cycle phase-specific and dynamic changes in the 3D structure of chromatin ^86^. Taken together, our single cell approach allowed us to make detailed predictions on sorting strategies, gene expression dynamics, fate couplings, growth factors implicated in fate control, and cell cycle modulation, which we validated to reveal novel fundamentals of early hematopoietic differentiation, as well as practical methods for further isolation and study of these cells.

## Acknowledgements

This work is funded by a Leukemia and Lymphoma Society Scholar award (1728-13), R01DK100915, R01099281 (MS). AMK is supported by a BW Fund CASI award and an Edward J Mallinckrodt Foundation Grant. SW and CW are supported by NIH training grant 5T32GM080177-07.

Data Availability: Data is available through an interactive tool (kleintools.hms.harvard.edu/paper_websites/tusi_et_al). The raw sequencing data and processed counts matrices were submitted to GEO (GSE89754).

## Methods

### Single-cell RNA-seq

#### Mice

For the basal state bone marrow sample (bBM), and for the sorted populations P1 to P5, bone marrow was harvested from 8-week old adult BALB/cJ female mice (Jackson Laboratories, Maine, USA). For the Epo-stimulated adult bone-marrow sample (eBM), 8-week old adult Balb/cJ female mice were injected with Epo (Procrit, Amgen corporation) sub-cutaneously once per 24 hours for a total of 48 hours, at 100 Units/25 g. For the fetal liver sample (FL), BALB/cJ female mice were set up for timed pregnancies, and fetal livers were harvested on embryonic day 13.5.

#### Cell preparation

*Tissue harvesting:* For bone-marrow preparation, femurs and tibiae were harvested immediately following euthanasia, and placed in cold (4°C) ‘staining buffer’ (phosphate-buffered saline (PBS) containing 0.2% Bovine Serum Albumin (BSA) and 0.08% Glucose). Bones were flushed using a 2 mL syringe with a 26-gauge needle and then crushed with a pestle and mortar to obtain all cells. Harvested bone marrow cells were filtered through a 40 μm strainer and washed in cold ‘Easy Sep’ buffer (PBS; 2% fetal bovine serum (FBS); 1mM EDTA). Fetal livers were prepared by mechanical dissociation in staining buffer and a wash in ‘Easy-Sep’ buffer.

*Positive selection for Kit+ cells:* bone marrow and fetal liver cell samples were each enriched for Kit expressing cells using magnetic beads, with the Mouse Biotin Selection Kit (STEMCELL technologies [Cat # 18556]) and Biotin Rat Anti-Mouse CD117 antibody (clone 2B8, BD Bioscience), following the manufacturer’s protocol.

*Density gradient centrifugation:* Following magnetic bead selection, dead cells and debris were removed from the bone marrow and fetal liver samples using density centrifugation in OptiPrep (Sigma, Cat # D1556). Briefly, cells were re-suspended in 0.5ml staining buffer, mixed with 1mL of 40% of Optiprep in PBS, and placed in a 5 mL tube. The cell suspension was carefully over-layered with 2 mL of 20% OptiPrep solution, and 1mL of 5% OptiPrep solution, and centrifuged at 800g for 15 min (Centrifuge break OFF). The top visible cell band that formed during centrifugation contained the live, Kit+ single cells, confirmed by flow cytometric analysis. This layer was carefully aspirated and used directly in the inDrops^1^ platform.

#### Single cell transcriptome droplet microfluidic barcoding using inDrops

For scRNA-seq, we used inDrops ^1^ following the protocol previously described ^2^ with the modifications summarized in **Extended Data Table 7**. Following droplet barcoding reverse transcription, emulsions were split into aliquots of approximately 1000 single cell transcriptomes and frozen at-80C. Two batches of Kit+ libraries were prepared, referred to as Batch 1 (bBM, n=840 cells; eBM, n=1, 141 cells; FL, n=1,953 cells) and Batch 2 (bBM, n=4,592 cells; eBM, n=1,314 cells; FL, n=7,529 cells) in **Extended Data Table 7**. These cell numbers correspond to the final number of transcriptomes detected upon sequencing (see “Cell filtering and normalization” below), and were in agreement with estimated inputs.

For the FACS subsets P1, P1-CD71^hi^, P2, P3, P4, and P5 (referred to collectively as “P1-P5”), all libraries were prepared in parallel, with a total of 16,206 cell barcodes detected in the sequencing data prior to filtering (P1, n=5,733 cells; P1-CD71^hi^, n=1,631 cells; P2, n=2,630 cells; P3, n=2,101 cells; P4, n=1,589 cells; P5, n=2,522 cells).

#### Sequencing and read mapping

The first batch of Kit+ (bBM, eBM, FL) libraries was sequenced on a HiSeq 2000, the remaining Kit+ libraries were sequenced on three NextSeq 500 runs, and all P1-P5 libraries were sequenced on a single NextSeq 500 run. Raw sequencing data (FASTQ files) were processed using the `inDrops.py` bioinformatics pipeline available at github.com/indrops/indrops and described in ^2^, with a few modifications. Bowtie version 1.1.1 was used with parameter −e 100; all ambiguously mapped reads were excluded from analysis; and reads were aligned to the Ensembl release 81 mouse mm10 cDNA reference.

#### Cell filtering and data normalization

Each sample (bBM, eBM, FL, P1-P5) was processed separately. The bBM, eBM, and FL samples (referred to collectively as “Kit+”) were initially filtered to include only abundant barcodes, based on visual inspection of the histograms of total reads per cell (see cell numbers reported in “Single cell transcriptome droplet microfluidic barcoding using inDrops”). An additional filtering step removed cells with transcript count totals in the bottom 5^th^ percentile (bBM, n=271 cells; eBM, n=148; FL, n=473). Subsets P1-P5 were filtered only by total transcript counts, with thresholds set by visual inspection of the total counts histograms (see cell numbers reported in “Single cell transcriptome droplet microfluidic barcoding using inDrops”). Next, we excluded putatively stressed or dying cells with >10% (bBM, eBM, FL) or >20% (P1-P5) of their transcripts coming from mitochondrial genes (bBM, n=165 cells; eBM, n=45; FL, n=698; P1, n=2,629; P1-CD71^hi^, n=879; P2, n=195; P3, n=69; P4, n=379; P5, n=62). Each cell’s gene expression counts were then normalized using a variant of total-count normalization that avoids distortion from very highly expressed genes. Specifically, we calculated 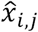, the normalized transcript counts for gene *j* in cell *i,* from the raw counts *x_i,j_* as follows: 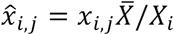, where *X* = Σ*_j_x_i,j_* and 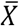 is the average of *Xi* over all cells. To prevent very highly expressed genes (e.g., hemoglobin) from correspondingly decreasing the relative expression of other genes, we excluded genes comprising >10% of any cell’s total counts when calculating 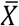 and *X_i_*.

### Exclusion of contaminating cell types and putative cell doublets

To clean up the data for the Kit+ samples, we clustered the single cell transcriptomes and excluded clusters that were identified as contaminating (non-HPC) cell types and putative cell doublets. No such clusters were detected in the P1-P5 samples. Clustering was performed as follows: we identified principal variable genes across the entire data set, as described in ^1^ i.e. genes that were highly variable (top 2000 most variable by v-score, a measure of above-Poisson noise [variability]), were expressed at non-negligible levels (at least 5 UMI-filtered mapped reads [UMIFM] in at least 3 cells), and which contributed to principal components with eigenvalues above those obtained following data randomization (n=59, n=35, n=71 principal components for bBM, eBM and FL samples, respectively). The expression level for each gene was standardized by a z-score transform (mean-subtraction, scaling by standard deviation), followed by density-based clustering (DBSCAN) ^3,4^ on a 2D PCA-tSNE plot (principal component analysis [PCA] followed by t-distributed stochastic neighbor embedding [tSNE^5^] as described in ^1^,^6^). The tSNE algorithm perplexity parameter was set to 30. Examination of marker gene expression in each cluster was then used to identify putative doublets and contaminating cell types.

In the bBM sample, two doublet clusters were identified: one co-expressed markers of mature macrophages and erythrocytes (n=38 cells), while the other co-expressed markers of granulocyte and erythroid progenitors (n=75 cells). The eBM sample included a cluster of mature macrophages (n=40 cells) but no identifiable cluster of doublets. The FL sample contained four contaminating cell types: vascular endothelium, hepatocytes, mesenchymal cells, and mature macrophages (n=769 cells total), in addition to a small cluster of doublets (n=18 cells). Doublets and contaminant cells were excluded from downstream analyses.

To increase confidence that putative doublet clusters were indeed combinations of two single cells, rather than true intermediate/transitional states, we generated simulated “artificial” doublets by randomly sampling and combining observed transcriptomes. We then applied PCA-tSNE clustering as described above to the union of observed and simulated cells, and identified clusters that were primarily composed of cells with a large number of doublet neighbors (two clusters in bBM, one in FL). These clusters were the same putative doublet clusters identified in the previous paragraph.

### Batch correction

Within each Kit+ sample, we observed batch effects between the first and second sequencing runs, with slightly fewer genes detected per cell in the second run compared to the first run. This was consistent with the choice of lower sequencing depth used in the second set of runs, but could also reflect differences in library preparation despite all cells being collected in a single droplet run. To prevent batch effects from distorting subsequent data analysis, for each sample we used the second (larger) batch to select variable genes and to calculate principal component (PC) gene loadings. Cells from all batches were then projected into the reduced space, and all subsequent analysis was performed on the reduced PC space.

#### Data visualization and construction of k-nearest neighbor (KNN) graphs

Following cell filtering, data was prepared for visualization and Population Balance Analysis (PBA) ^7^ by constructing a kNN graph, in which cells correspond to graph nodes and edges connect cells to their nearest neighbors. A kNN graph was constructed separately for each of the three Kit+ samples and for the merged P1-P5 samples (note that the kNN graph for P1-P5 was used only for the visualization in **Extended Data Fig. 5**).

For the Kit+ samples, genes with mean expression >0.05 and coefficient of variation (standard deviation/mean) >2 were used to perform principal components analysis (PCA) down to 60 dimensions (bBM, eBM, FL). For all analyses in this paper, data were z-score normalized at the gene level prior to PCA (qualitatively similar results were also obtained without z-score normalization, which weights highly expressed genes more heavily than lowly expressed genes). After PCA, a kNN graph (k=5) was constructed by connecting each cell to its five nearest neighbors (using Euclidean distance in the PC space).

For P1-P5, highly variable genes were filtered using the v-score statistic (above-Poisson noise) rather than CV, keeping the top 25% most highly variable genes and requiring least 3 UMIFM detected in at least 3 cells (n=3,459 genes). Additionally, a strong cell cycle signature was observed in the initial graph visualization, manifested by colocalization of cells expressing G2/M genes (Ube2c, Hmgb2, Hmgn2, Tubal1b, Mki67, Ccnb1, Tubb, Top2a, Tubb4b). Therefore, we constructed a G2/M signature score by summing the average z-score of these genes, then removed genes highly correlated (Pearson r > 0.2) with the signature (n=31 genes). Finally, the kNN graph was constructed with k=4 using the first 30 PCs.

The kNN graphs were visualized using a force-directed layout using a custom interactive software interface called SPRING^8^. For the Kit+ samples, several manual steps were taken to improve visualization. It is important to emphasize that the manipulations affect visualization only. All subsequent analyses depend on the graph adjacency matrix, which is not affected by any of the changes to the graph layout. For visualization purposes, we manually extended the length of the Mk, Ba, and Mo branches by pinning the position of cells at the end of each branch, and allowing the remaining structure to follow. In the bBM sample, we compressed the CEP “bulge” region of the graph by bringing its bounding cells together.

#### Smoothing over the kNN graph

We smoothed data over the kNN graph for gene expression visualization and for one analysis (“Global changes in gene expression in stress conditions”). Smoothing was done by diffusing the property of interest (e.g., gene expression counts or number of mapped cells) over the graph, as described in ^9^. In brief, let *A* be the adjacency matrix of the kNN graph, where *A_i,j_*=1 if an edge in the graph connects nodes *i*and *j.* Define *A*\*as the transition matrix, obtained by row-normalizing *A*:

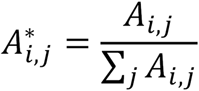

Let *E_i_* be the quantity of interest (e.g., expression level) in cell *i.* Then *E*,* the smoothed vector of *E*, is computed as follows:

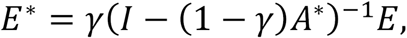

where *γ* is a diffusion constant (*γ* = 0.05 in all presented analyses) and *I* is the identity matrix.

#### Formal measure of the continuity of transcriptional states

To demonstrate that the continuum appearance of the Kit+ transcriptomes was not a trivial outcome of our analysis methods, we used the same tools to analyze an scRNA-seq dataset of mature blood cells (peripheral blood mononuclear cells, PBMCs) [https://support.10xgenomics.com/single-cell-gene-expression/datasets/2.0.1/pbmc8k], which consist of several distinct cell types (**Extended Data Fig. 1a**). In addition to generating a SPRING plot of the data, we also assessed each dataset’s interconnectivity by examining the behavior of random walks over the kNN graphs, as in Velten et al.^10^. In detail, after subsampling the PBMC data to contain the same number of cells as the bBM dataset, we applied PCA and constructed a kNN graph (k=10) for each dataset. We then simulated 1,000 random walks for each graph and plotted the fraction of nodes (cells) visited as a function of the number of steps (**Extended Data Fig. 1a**).

#### Population balance analysis (PBA)

The PBA algorithm calculates for each cell a scalar “potential” that is analogous to a distance, or pseudotime, from an undifferentiated source, and a vector of fate probabilities that indicate the distance to fate branch points. These fate probabilities and temporal ordering were computed using the python implementation of PBA (available online https://github.com/AllonKleinLab/PBA), as described in Weinreb et al. ^7^.

The inputs to the PBA scripts are a set of csv files encoding: the edge list of a k-nearest neighbor graph of the cell transcriptomes (A.csv); a vector assigning a net source/sink rate to each graph node (R.csv); and a lineage-specific binary matrix identifying the subset of graph nodes that reside at the tips of branches (S.csv). These files are provided in the **Supplementary Data** for the BM and FL data sets. PBA is then run according to the following steps:

1. Apply the script “`compute_Linv.py −e A.csv`”, here inputting edges (flag “−e”) from the SPRING kNN graph (see previous Methods section). This step outputs the random-walk graph Laplacian, `Linv.npy`.
2. Apply the script “`compute_potential.py –L Linv.npy –R R.csv`”, here inputting the inverse graph Laplacian (flag “–L”) computed in step (1) and the net source/sink rate to each graph node (flag “–R”). This step yields a potential vector, `V.npy`, that is used for temporal ordering (cells ordered from high to low potential). The vector R provided in the **Supplementary Data** was estimated as described in the next section.
3. Apply the script “`compute_fate_probabilities.py –S S.csv –V V.npy –e A.csv –D 1`”, here inputting the lineage-specific exit rate matrix (flag “-S”), the potential (flag “–V”) computed in step (2), the same edges (flag “-e”) used in step (1) and a diffusion constant (flat “–D”) 1. This step yields fate probabilities for each cell.

Front end **Figures 1–6** make use of PBA analyses of bBM data. For **Fig. 4e** and **Extended Data Fig. 11**, a temporal ordering of erythroid differentiation was generated for the FL data set using the same steps, with input files also provided in **Supplementary Data**.

#### Estimation of net source/sink rate vector R

*Theory:* A complete definition of the vector *R* in terms of biophysical quantities is provided in Weinreb et al. ^7^. In brief, for a gene expression space described by a vector *X=(X_1_,X_2_,…,X_N_*) giving the expression of each of *N* genes, *R(x)* gives the net imbalance between cell division and cell loss locally for cells with gene expression profile x. *R(x)* is corrected for cell enrichment and loss resulting from experimental procedures such as sample enrichment, as follows. In this experiment, all progenitors including HSCs express Kit, but eventually down-regulate it as they terminally differentiate. Thus, no cells enter the experimental system other than through proliferation of existing Kit+ HPCs, but the selection for *Kit+* cells during sample isolation induces a net sink on cells down-regulating Kit expression. For a self-renewing system, cell division and cell loss are precisely balanced, so ∫*R(x)dx* = 0. To apply PBA, one does not need to estimate *R(x),* but only its value at points x_i_ at which the *M* cells *i*=1,…,*M* are observed in the scRNA-Seq measurement. Thus *R* is a vector over the cells in the system. For a self-renewing system, the sum over all cells satisfies the same constraint, Σ*_i_ R_i_* = 0.

*Estimation of R:* We assigned negative values to *R* for the top 10 cells with highest marker gene expression for each of the seven terminal lineages (see **Extended Data Table 1** for marker genes), which were separately confirmed to show reduced Kit expression. We assigned different exit rates to each of the seven lineages using a fitting procedure that ensured that cells identified as putative HSCs would have a uniform probability to become each fate. Putative HSCs were identified by the similarity of their transcriptomes to microarray profiles from the ImmGen database (we used SC.LT34F.BM [long-term BM HSCs] for bBM and SC.STSL.FL [short-term FL HSCs] for FL; for more details, see section “ImmGen Bayesian classifier” below). We assigned a single positive value to all remaining cells, with the value chosen to enforce the steady-state condition Σ*_i_R_i_* = 0. In the fitting procedure, all exit rates are initially set to 1 and iteratively incremented or decremented until the average fate probabilities of the putative HSCs were within 1% of uniform. The resulting vector *R* is provided in the **Supplementary Data** file. The separate lineage exit rates were then used to form the lineage-specific exit rate matrix S, provided in the **Supplementary Data**.

#### Assignment of PBA fate probabilities and temporal ordering to eBM dataset

We assigned to each of the eBM cells the average temporal order (or potential *V*) and average fate probabilities of the 20 mostly similar basal BM cells. To do this, we first carried out a principal component analysis (PCA) on the basal BM cells into 60 dimensions. We then used the gene loadings of the 60 PCs to project the eBM data into the same PC space. The distance of each eBM cell to each basal BM neighbors was then measured by cosine distance in the 60-dimensional sub-space.

#### ImmGen Bayesian classifier

We used a published microarray profile^11^ to search for similar cells in our own dataset using a naïve Bayesian classifier, implemented as follows.

The Bayesian classifier assigns cells to microarray profiles based on the Likelihood of each microarray profile for each cell, with the Likelihood calculated by assuming that individual mRNA molecules in each cell are multinomially sampled with the probability of each gene proportional to the microarray expression value for that gene. Consider a matrix *E* of mRNA counts (UMIs) with *n* rows (for cells) and > columns (for genes), and also a matrix *M* with *m* rows (for microarray profiles) and > columns for genes. *M* was quantile normalized and then each microarray profile was normalized to sum to one. *E* was normalized as in section “Cell filtering and data normalization”. The (*n* × *m*) matrix *S_ij_* giving the Likelihood of each microarray profile *j* for each cell *i* is,

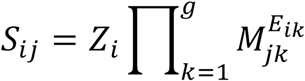

where *Z_i_* is a normalization constant that ensures Σ*_i_S_ij_* = 1.

#### Computing the hematopoietic lineage tree

We used the fate probabilities from PBA to infer the topology of the hematopoietic lineage tree using an iterative approach (**Fig. 1e-f**). Each iteration began with a set of fates and a probability distribution over those fates for each cell. For every pair of fates, we computed a fate coupling score (see next paragraph) and merged pairs with a score significantly higher than expected under a null model. The merged fates inherited probabilities from the starting fates by simple pairwise addition.

The coupling score between two fates *A* and *B* is the number of cells with *P(A)P(B)* > *ε*, where we used a value *ε* = 1/14 throughout. To generate a null distribution for each fate pair, we computed pairwise coupling scores for 1000 permutations of the original fate probabilities. The heatmaps in **Fig. 1e** show z-scores with respect to these null distributions.

#### Analysis of fate-correlated genes at hematopoietic choice-points

To discover fate-associated genes at key choice points in hematopoiesis (**Extended Data Fig. 2; Extended Data Table 2**), we ranked transcription factors (TFs) and cell surface markers (TFs from http://genome.gsc.riken.jp/TFdb/tf_list.html, surface markers from http://www.ebioscience.com/resources/mouse-cd-chart.htm) by their correlation with PBA-predicted fate probability, restricting to cells that were bi-potent for the given choice. Specifically, to find TFs associated with fate *A* at an *A/B* choice point, we first selected cells with *P(A) * P(B) > ε*, (*ε*=1/14) and then ranked the TFs by their correlation with the fate bias [*P(A)*-*P(B)*]. In **Extended Data Table 2**, we report all genes with Pearson significance *p* < 0.01 (Bonferroni corrected). In **Extended Data Fig. 2**, we show at most 10 genes for any one choice point.

#### Mapping P1-P5 subsets to the Kit+ graphs

For **Fig. 2b**, cells from subsets P1-P5 were projected into the same PC space as the bBM data, then mapped to their most similar Kit+ neighbors. In detail, first, counts were converted to transcripts per million (TPM) for all samples. Then, using only the bBM cells, the 3000 most highly variable genes (measured by v-score) with at least 3 UMIFM in at 3 cells were z-score normalized and used to find the top 50 principal components. Next, the P1-P5 subset cells were z-score normalized using the gene means and standard deviations from the bBM data and transformed into the bBM PC space. Lastly, each P1-P5 cell was mapped to it closest bBM neighbor in PC space (Euclidean distance).

#### Extracting MPP-to-Erythroid trajectory cells

To isolate the erythroid trajectory, we defined an MPP-to-erythroid axis in each of the three Kit+ datasets by ordering cells based on their graph distance from unbiased MPPs (cells identified based on the ImmGen classifier as described above), and keeping only cells for which the probability of erythroid fate increased or remained constant with graph distance. Graph distance was measured by PBA potential, and starting with the cell closest to the HSC origin, we added the cell with next-highest potential to the trajectory if the PBA-predicted erythroid probability for cell *i* was at least 95% of the average erythroid probability of the cell(s) already in the trajectory.

More formally the procedure is as follows: order all *N* cells in the experiment from highest to lowest PBA potential *V,* with decreasing potential corresponding to increasing distance from MPPs ^7^. Let *E_i_* be an indicator variable for the membership of ordered cell *t_i_* in the erythroid trajectory (*E_i_* = 1 if cell *i* is in the trajectory; otherwise, *E_i_* = 0). If *P_i_* is the PBA-predicted erythroid probability for ordered cell *i,* then *E_i_* = 1 if

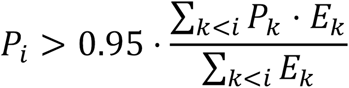

Erythroid trajectory cells were then ordered by decreasing potential. Defining *t_i_* as the index of the *j^th^* erythroid trajectory cell,

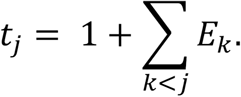

Throughout the paper, we report this cell order (akin to the “pseudotime” reported in other publications) as a percent of ordered cells, with the first, least differentiated cell at 0% and the most mature cell at 100%. This is not meant to suggest that erythroid differentiation ends with this final observed cell.

#### Identifying dynamically varying genes

For each gene, a sliding window (n=100 cells) across the MPP-to-Erythroid ordering was used to identify the windows with maximum and minimum average expression as in ^6^. A t-test was then performed to assess the statistical significance of the difference in expression levels. To estimate the false-discovery rate (FDR), we permuted the order of the cells and repeated the above analysis^6^. For a p-value generated by the observed (non-permuted) ordering, the FDR-corrected p-value is the fraction of genes from the permuted ordering with that p-value or less. Any gene with an FDR-corrected p-value <0.05 was considered significantly variable.

#### Identifying stage transitions in MPP-to-Erythroid trajectory

Transition points between stages of erythropoiesis were defined using the frequency of gene inflection points (**Fig. 4b**), patterns of PBA-predicted fate probabilities (**Fig. 1c**), and the fate potentials of FACS subsets P1-P5 (**Figs. 2,3**). That said, due to the continuous nature of the transcriptional states, the locations of these transitions should be considered approximate.

The inflection point density is the number of genes turning on or off at a given point on the trajectory. For each gene, inflection points were identified as the points with maximally increasing or decreasing expression as follows: first, each dynamically varying gene’s trajectory was smoothed using Gaussian smoothing with a width o=5% of total trajectory. The gene expression derivative for gene k, denoted 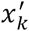 was then computed by taking a 10-cell moving average of the difference between consecutive smoothed gene expression values. Inflection points were then identified as the points with maximum or minimum derivative for each gene. To exclude maxima or minima resulting from relatively small gene expression fluctuations, only appreciably large extrema were kept for further analysis. Specifically, the point with the maximum derivative for gene *k,* max (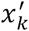), was kept only if

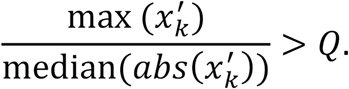

Minima were similarly filtered, requiring the ratio to be <-*Q*. We chose a threshold 0=6, but results do not qualitatively change over a range of *Q*. We then plotted the density of these inflection points over the MPP-to-erythroid axis. Regions with large-scale gene expression changes have a high density of inflection points, while a low density characterizes relatively stable states.

#### Dynamic gene clustering

Dynamically varying genes were clustered based on their behavior at the transition points. To prevent overfitting, we used only three transitions (3%, 18%, 86%) by splitting the EEP state and assigning the first and second halves to the EBMP and CEP states respectively. At each transition, genes were classified as increasing, decreasing, or unchanging, giving a total of 3^3^=27 possible patterns. After smoothing gene expression traces, the data were binned by calculating the mean expression in each of the four stages. To remove noisy genes or genes that varied little across bins, we calculated the range of binned expression values, range(*x_i,binned_*} = ^max^(*x_i,binned_*} − min (*x^i,binned^*), for each gene and proceeded with the top 50% most variable genes. Next, to place all genes on a similar scale, each gene’s binned expression values were divided by its maximum binned value. Finally, the differences between consecutive bins were thresholded: differences > 0.15 were called increasing, differences <-0.15 were called decreasing, and differences >-0.15, < 0.15 were called unchanging.

#### Gene set enrichment analysis (GSEA)

Each of the 27 gene clusters was used as input for GSEA (Hypergeometric test), using all genes as background. Ribosomal genes were excluded from the input, as were predicted genes (Gm*). Gene sets from MSigDB v5.1^12^ from the following list were tested for enrichment: Hallmark (h.all.v5.1.symbols.gmt), C2 curated canonical pathways (c2.cp.v5.1.symbols.gmt), C3 transcription factor targets (c3.tft.v5.1.symbols.gmt), and C5 Gene Ontology (c5.all.v5.1.symbols.gmt). Additionally, for transcription factor target (TFT) enrichment analysis, we used gene sets from the ChEA database^13^.

#### Cell cycle phase analysis

Genes with periodic expression correlated with the cell cycle in HeLa cells ^14^ were used to generate a cell cycle phase score for each cell. The list of phase-specific genes was filtered to exclude genes with a mean expression >25 TPM in MPP-to-erythroid trajectory cells. For **Fig. 6a**, a sliding window average was computed using a window size of 10% MPP-to-erythroid progression (~200 cells) and a jump size of 5%. For **Fig. 6b**, counts were normalized by the mean expression at the gene level, and smoothed using Gaussian smoothing. Then, for each phase (G1/S, S, G2/M, M, M/G1), a phase score was calculated by averaging the smoothed gene expression traces for the genes specific to that phase.

#### Testing the influence of cell cycle genes on MPP to Erythroid cell order

To test the extent to which cell cycle genes influenced the ordering of cells along the MPP to Erythroid trajectory, we excluded annotated cell cycle genes (as in ^15^)-a combination of genes from the Gene Ontology database (G0:0007049) and Cyclebase^16^-and repeated kNN graph construction and PBA. As shown in **Extended Data Fig. 13**, the resulting cell order was largely unchanged, as were the dynamics of cell cycle genes.

#### Identifying genes that change steadily in the CEP stage

To identify genes that are steadily up-or down-regulated throughout the CEP (**Fig. 6c** and **Extended Data Table 6**), we tested each gene’s magnitude of change (slope) and the linearity of its change (the error of the actual gene trace from a straight line). Restricting to cells in the CEP stage and genes with at least 2 UMIFM in at least 5 cells, we fit a linear regression to the ordered gene expression values and also generated a smoothed expression trace using a Gaussian kernel (width σ=5%). We then computed a “linearity score” for each gene by dividing the slope of the regression line by the root-mean-square error between the regression line and smoothed trace. Steadily increasing genes receive large positive scores, while steadily decreasing genes are assigned a large negative score. Genes that do not change much or that change non-linearly (e.g., sharply increasing only at the end of the stage) receive scores close to 0.

#### Global changes in gene expression in stress conditions

Cells from eBM and FL (stress samples) were mapped to their most similar bBM counterparts, and differentially expressed genes were identified. Mapping was carried out by applying PCA to the bBM and stress samples and finding the closest 20 bBM neighbors for each stress cell. Specifically, the input genes were the principal variable genes described in the “Cell filtering and data normalization” section. Counts matrices were z-score normalized separately for each sample, and PCA was performed on the basal sample to obtain the gene loadings. Using the top 60 PCs, each sample was then transformed using these coefficients, thereby projecting the cells into the same PCA space. To validate this mapping method, we performed the same procedure using different subsets of bBM data as training and test sets (see “Validation of cross-sample cell mapping” below)

Each stress cell’s 20 closest bBM neighbors (Euclidean distance) were found, and for the purpose of comparing gene expression, each of these *k* (20) neighbors inherited 1/k (1/20) of the transcript counts from the mapped stress cell. To enable the comparison of regions of gene expression space (as opposed to comparing single mapped cells to single basal cells), the mapped and original gene expression values were smoothed over the kNN graph, as described in a previous section (“Smoothing over the kNN graph”). To avoid comparing gene expression patterns in regions that were relatively unpopulated in the stress sample (e.g., parts of the granulocyte branch), we smoothed the number of mapped stress cells per basal cell over the graph and then excluded basal cells with few mapped stress cells (number mapped cells <=9 for eBM and >=20 for FL).

A differential expression score for each cell *i* and gene *j* was defined as the max-normalized difference between mapped and basal expression, 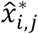 and 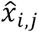, respectively:

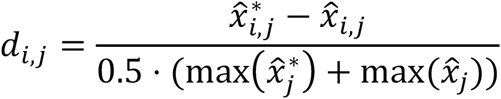

A gene level score, *D_j_*, was created by summing over the cells, *D_j_* = Σ*_i_d_i,j_*. Genes were considered differentially expressed if

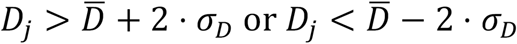

where 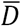 is the average over all gene level scores *D_j_* and σ*_D_* is the standard deviation.

Then, for each differentially expressed gene, the gene was counted as differentially expressed at a given cell if

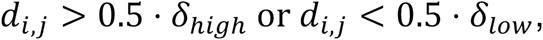

where *δ_high_* is the 99^th^ percentile of *D_j_* and *δ_low_* is the 1^st^ percentile of *D_j_*.

#### Validation of cross-sample cell mapping

To test the accuracy of the method for mapping eBM and FL cells to bBM cells, we divided the bBM sample into a training set (random sample of 75% of the cells) and test set (the remaining 25%). The mapping procedure described in the previous section was then used to map the test set to the training set. As one measure of the accuracy of the mapping, we assigned the test cells the average PBA-predicted fate probabilities and differentiation ordering of the training cells to which they mapped. Both measures were relatively unchanged from their original values (Spearman correlation of 0.97 for the differentiation ordering and >0.95 for each fate probability). As a second measure, we repeated the test for finding global changes in gene expression, using the same gene level score *(Dj)* cutoff as for the eBM. This revealed no significantly differentially expressed genes between the training and test sets.

#### Region-specific differential expression

Prior to identifying differentially expressed genes, we excluded genes with large batch effects. While different sequencing depths led to a small change in the average expression of many genes from the first batch to the second, a small number of genes showed major batch effects beyond this, presumably due to differences in library prep. We performed a binomial test for differential expression ^17^ between the two batches of cells and excluded genes with *p*<10^−50^, resulting in the removal of 461 genes.

In general, genes can be differentially expressed globally or only in specific cell populations. Particularly when comparing FL to bBM, many genes showed global up-or down-regulation. In order to identify differentially expressed genes likely to important specifically for erythropoiesis (or in a particular stage of erythropoiesis), we created a region-specific differential expression score, described in detail below. This score measures the magnitude of the expression difference within a region of interest (ROI) relative to the magnitude outside the region; genes with a larger difference within the ROI than outside of it receive a high score (positive for up-regulation, negative for down-regulation). For the analyses in this paper, we tested for differential expression in five ROIs: the erythroid trajectory stages EBMP, EEP, CEP, and ETD, plus an expanded selection of “MPP” cells, which included cells with a maximum PBA-predicted lineage probability (for all lineages) <0.4, excepting cells already included in one of the erythroid trajectory stages.

After mapping stress cells to their single closest neighbor in bBM (as described in the previous section), we selected bBM cells in the ROI and the stress cells mapping to them. We first identified genes differentially expressed within the ROI by performing a binomial test for differential expression^17^, which tests the probability that a gene is expressed more frequently in one population than another. After correcting for multiple hypothesis testing (Benjamini-Hochberg procedure^18^), we proceeded with genes with an FDR-corrected p-value <0.05.

To identify genes differentially expressed *specifically* within the ROI and not elsewhere, we calculated the mean-normalized expression difference for ROI cells and non-ROI cells for the significant binomial test genes. For two samples, *A* (stress) and *B* (basal), the mean-normalized expression difference of gene *i*within the ROI, *y_in,i_*, is

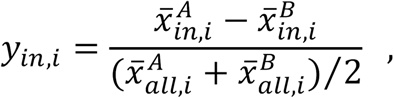

where 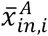 is the average expression of gene *i* within the ROI in sample A. A similar score was calculated for cells outside the ROI:

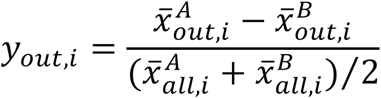

Plotting *y_in,i_* vs. *y_out,i_* clearly reveals genes more highly DE within the ROI than without. A single score per gene was computed as follows:

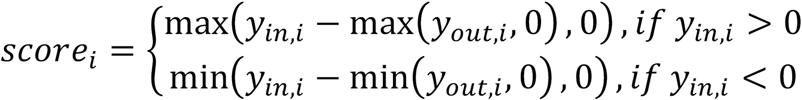

Intuitively, this score is large and positive if a gene is more strongly upregulated within the ROI than without, is large and negative if a gene is more strongly downregulated within the ROI than without, and is close to 0 otherwise.

To build gene lists for GSEA input, we first selected genes with *score_t_ >* 0.1 * max *(score)* (for upregulated genes) or *score_i_ <* 0.1 * min *(score)* (for downregulated genes) and then used the top 100 genes by binomial test p-value.

#### Flow cytometric sorting for P1 to P5 subsets

Bone marrow cells from adult BALB/cJ male or female mice, ages 8-12 weeks, were lineage-depleted using the Mouse Streptavidin RapidSpheres Isolation Kit (STEMCELL Technologies [Cat# 19860A]), with the following biotinylated antibodies:

anti-CD11b (Clone M1/70 [#557395], BD Biosciences)

anti-Ly-6G and Ly-6C (Clone RB6-8C5 [#553125], BD Biosciences)

anti-CD4 (Clone RM4-5 [#553045], BD Biosciences)

anti-CD8a (Ly-2) (Clone 53-6.7 [#553029], BD Bioscience)

anti-CD19 (Clone 1D3 [#553784], BD Biosciences)

anti-TER119 (Clone TER119 [#553672], BD Biosciences).

Lineage-depleted cells were then labeled with the following antibodies in the presence of 1% rat serum:

streptavidin Alexa Fluor 488 (Molecular Probes), to mark lineage-positive cells

CD117-APC Cy7 (Clone 2B8 [#105826], Biolegend)

TER119-BUV395 (Clone TER-119 [#563827], BD Biosciences)

CD71-PE Cy7 (Clone RI7217 [#113812], Biolegend)

CD55-AF647 (Clone RIKO-3 [#131806], Biolegend)

CD105-PE (Clone MJ7/18 [#120408], Biolegend)

CD150-BV650 (Clone TC15-12F12.2 [#115931], Biolgened)

CD41-BV605 (Clone MWReg30 [#133921], Biolegend)

CD49f (=itga6)-BV421 (Clone GoH3 [#313624], Biolegend)

Following washes, cells were re-suspended in DAPI-containing buffer and sorting was performed on BD FACSAria II with a 100 n nozzle. Sorted populations were defined as in **Fig. 2a**.

#### qRT-PCR on sorted populations

RNA was prepared from sorted cell subsets using the RNeasy Micro Kit (QIAGEN; CAT# 74004) or TRIzol reagent (Ambion; CAT# 15596026), and measured with RiboGreen RNA reagent kit (Thermo Scientific) on the 3300 NanoDrop Fluorospectrometer. cDNA was synthesized using the same amount of input RNA for all samples in a parallel reaction, using the Super Script III first-strand synthesis system for RT-PCR (Invitrogen) with random hexamer primers. The ABI 7300 sequence detection system, TaqMan reagents and TagMan MGB probes (Applied Biosystems, San Diego, CA) were used following the manufacturer’s instructions. qPCR was carried on 4 serial dilutions of each cDNA sample, and the linear part of the template dilution/signal response curve was used to calculate relative mRNA concentrations following normalization to β-actin, using the ΔCt method.

The following TagMan MGB probes were used:

Mst1r (Mm00436382_m1), Ryk (Mm01238551_m1), IL17ra (Mm00434214_m1), Mt2 (Mm00809556_s1), Slc26a1 (Mm01198850_m1), Slc4a1 (Mm00441492_m1), Trib2 (Mm00454876_m1), Cd34 (Mm00519283_m1), Meis1 (Mm00487664_m1), Hpn (Mm01152654_m1), Pf4 (Mm00451315_g1), Dntt, (Mm00493500_m1), Ms4a2 (Mm00442778_m1), Elane (Mm00469310_m1), S100a9 (Mm00656925_m1), F13a1 (Mm00472334_m1), Egr1(Mm00656724_m1), Apoe (Mm01307193_g1), Ldb1 (Mm00440156_m1), Zfpm1 (Mm00494336_m1), Tfrc (Mm00441941_m1), Hbb-b1 (Mm01611268_g1), Alas2 (Mm01260713_m1), Band3 (Mm01245920_g1), Nfe2 (Mm00801891_m1), Gata1 (Mm01352636_m1), Gata2 (Mm00492300_m1), Klf1 (Mm00516096_m1) and PU1/Sfpi1 (Mm00488393_m1).

#### Colony-formation assays in methylcellulose for P1 to P5 and Kit+CD55^-^ cells

From each freshly sorted cell population, 10,000 cells were mixed with 1 ml MethoCult (M3234, STEMCELL Technologies) supplemented with EPO (2U/ ml), SCF (50ng /mL), IL-310 (ng/ mL) and IL-6 (10 ng/mL). Erythroid (CFU-e or BFU-e) and GM colonies were scored from triplicate plates on days 3, 4 and 7 of culture. Hemoglobin expression in erythroid colonies was verified by staining with diaminobenzidine in situ before scoring.

For megakaryocyte, colony formation assay was carried out using MegaCult®-C Complete Kit (Catalog #04970/04972) with added TPO (50 ng/ mL), IL-3 (10 ng/ mL), IL-6 (20 ng/ mL) and IL-11 (50 ng/ mL). From each freshly sorted subset, 10,000 cells were plated in double chamber slides. On day 7 of culture, the slides were dehydrated, fixed in ice-cold acetone, and stained for acetylcholinesterase.

#### Bulk Liquid cultures of sorted cell populations

Sorted cells were cultured in Iscove’s Modified Dulbecco’s Medium (IMDM) in the presence of 20% FCS supplemented with SCF (50 ng/ mL), IL-3 (10 ng/ mL), IL-6 (10 ng/ mL), EPO (2 U/ mL), TPO (50 ng/ mL), IL-11 (50 ng/ mL) and IL-5 (10 ng/ mL) for 7 days. Cells were collected on days 2, 5 and 7, and labeled with the following cell surface markers for flow cytometric analysis:

TER119-BV421 (Clone TER-119, #116233 Biolegend)

CD71-PE Cy7 (Clone RI7217, #113812] Biolegend)

CD117-APC Cy7 (Clone 2B8, #105826 Biolegend)

FcsRIa-AF700 (Clone MAR-1,#134323 Biolegend)

CD41-BV605 (Clone MWReg30, #133921 Biolegend)

Cd11b-PE Cy5 (Clone M1/70, #101209 Biolegend)

Ly 6G/C-FITC (Clone RB6-8C5, #553126 BD Biosciences).

#### Single cell liquid cultures of mouse BM progenitors

Freshly harvested mouse bone-marrow was labeled with the same antibody scheme as

detailed above, to allow identification of the Kit+ gates for P1 to P5 and CD55^-^. Single

cells were sorted from each of these gates into 96-well plates, retaining index-sorting

parameter for each cell, using a BD FACSAria II with a 130 n nozzle. Cells were

cultured for 3 to 10 days, in IMDM+ 20% FBS, with the following added growth factors:

SCF (50 ng/mL): Recombinant Murine SCF, #250-03 Peprotech

IL-3 (10 ng/mL): Recombinant Murine IL-3, #213-13 Peprotech

IL-6 (10 ng/mL): Recombinant Murine IL-6, #216-16 Peprotech

EPO (2U/mL): PROCRIT® (epoetin alfa), #606-10-971-8

IL-11 (50 ng/mL): Recombinant Murine IL-11, #220-11 Peprotech

IL-5 (10ng/mL): Recombinant Murine IL-5, #215-15 Peprotech

TPO (50 ng/mL): Recombinant Murine TPO, #315-14 Peprotech

G-CSF (15 ng/mL): Recombinant Murine G-CSF, #250-05 Peprotech

GM-CSF (15 ng/mL): Recombinant Murine GM-CSF, #315-03 Peprotech

Fresh growth factors were added to the medium of each well on days 4 and 8. The clones in each well were labeled on days 3, 7 or 10, with the same antibody cocktail as detailed above under “Bulk liquid cultures”, but with concentration for each antibody batch that were first optimized with appropriate titrations, to minimize non-specific binding under conditions of low cell number. Clones were analyzed using the High Throughput Sampler (HTS) attachment of the BD LSR II.

#### Fate co-occurrence from single cell liquid culture data

To measure the significance of fate co-occurrence from the single-cell fate assay data, we employed a method similar to that described for calculating fate couplings from the PBA predictions (methods section “Computing the hematopoietic lineage tree”). Since we assayed clonal fate from each FACS subset separately, clones were not represented at the same frequency as in the Kit+ pool (number of clones assayed: CD55^-^, n=58; P1, n=287; P2, n=324; P3, n=125; P4, n=96; P5, n=268; average frequency in Kit+ population: 59.1% CD55^-^, 21.4% P1, 6.6% P2, 4.0% P3, 0.8% P4, 4.9% P5). To adjust for this, we randomly resampled the clone data to ensure clones from each subset were represented in the same proportion as in the Kit+ population (originally: n=1,158 clones; after resampling: n=8,000 clones). We then computed the observed fate co-occurrence for each fate pair as the number of clones with >2% of cells of the two fates (permitting the presence of other fates as well). Next, we estimated the null distribution by shuffling each fate’s data separately (2000 replicates) and counting fate co-occurrence as above. Lastly, we calculated the significance of each fate pair’s co-occurrence as the z-score of the observed co-occurrence with respect to the null distribution.

#### Growth factor perturbations of erythroid colony formation

CFU-e and BFU-e colony formation assays in MethoCult (M3234 STEMCELL Technologies) were carried out on either freshly isolated bone marrow or on embryonic day 13.5 fetal liver cells from Balb/cJ mice. The following growth factors were tested: MSP/MST1 (R&D systems; CAT #6244-MS-025), Recombinant Human/Mouse Wnt-5a (R&D systems; CAT #645-WN-010) and Recombinant Murine IL17 (IL-17A) (PeproTech; CAT #210-17). In each experiment, a range of Epo concentrations was tested, with or without added additional growth factors (MSP, Wnt5a or IL-17A) as indicated in **Fig. 5** and in **Extended Data Fig. 12**. In addition to Epo, IL-3 (10ng/mL) and SCF (50ng/ mL) were added to the MethoCult in the case of BFU-e assays. Each condition was tested in quadruplicates, in at least 2 separate experiments. Colonies were scored on day 3 (for CFU-e), day 4 (for late BFU-e) and on day 7 (for early BFU-e) following staining with diaminobenzidine, to highlight hemoglobin expression.

**IL17RA-deleted mice:** To generate the IL-17RA-deleted line, IL-17RA flox/+ mice^19^ were bred with CMV-Cre mice (#003465, JAX lab). The generation of *il17ra del* allele in the F1 generation of il17raflox/+ x CMV-Cre mating pairs were screened by PCR of tail DNA. To remove the CMV-Cre allele present in the F1 generation, IL-17RA *del/+;* CMV-Cre+/-mice were outcrossed with B6 mice.

### Colony-formation assays with human bone-marrow

Human bone-marrow mononuclear cells (MNCs) (85,000 cells, STEMCELL Technologies 70001.1) were mixed with 1 ml MethoCult (STEMCELL Technologies H4230) supplemented with either EPO (0.05U/ ml), and in the presence or absence of IL-17a (R&D systems 7955-IL-025). CFUe colonies were scored from triplicate plates on day 7.

#### Cell cycle studies

##### Flow cytometric cell cycle analysis of bone-marrow cells*in vivo*

These were carried out as described^20^. Briefly, BrdU (100 μl of 10 mg/ml stock in PBS) was injected intra-peritoneally into adult mice 30 minutes prior to euthanasia. Following bone marrow harvesting, cells were immediately placed in cold staining buffer and labeled with LIVE/DEAD kit (Invitrogen) to identify dead cells, and were then fixed and permeabilized. Cell surface staining for each of the 5 subsets P1 to P5 was carried out as described above. Simultaneously, DNA-incorporated BrdU was detected using a biotin-conjugated anti-BrdU antibody (Abcam) following mild digestion with DNaseI. DNA content was assayed by labeling with 7AAD (BD Biosciences). Cells were then analyzed for cell surface labeling, BrdU incorporation and DNA content by flow cytometry.

##### Cell cycle arrest studies during erythroid differentiation *in vitro*

BM cells were harvested and immediately enriched for Kit+Lin^-^ TER119^-^ CD71^-^ cells using magnetic beads, as described above. The enriched cell fraction was placed in culture in IMDM/ 20% FCS/ Epo (2 U/ mL) at time 0, in the presence or absence of aphidicolin (6μM, Sigma CAT#A0781). At t= 10 hours, all the cells were washed 3 times in culture medium to remove aphidicolin, and returned to culture, which continued for up to a total of 36 hours.

At the indicated time points, cell aliquots were taken for: RNA extraction and qRT-PCR for Hbb-b1 and b-actin; and for a simultaneous flow-cytometric analysis of CD71, Ter119 expression and cell cycle status. For the latter, cells were pulsed in vitro for 25 minutes prior to collection with BrdU (33 μM), then processed as described above for BrdU incorporation, DNA content and cell surface CD71/Ter119 expression.

##### Western blot analysis

*Cells:* Bone marrow cells were sorted as above, except that the P1 population was further subdivided into CD71^med^ and CD71^hi^ subsets. For negative controls, we used 3T3-L1 cells. For positive controls, 3T3-L1 cells were transduced with the MICD4-GATA1 retrovirus as described^20^. Cell pellets were snap-frozen in liquid nitrogen following the sort.

Cell lysates were quantified by the BCA Protein Assay Kit (Pierce) and separated by SDS-PAGE gel electrophoresis. PVDF membranes were probed with antibodies against GATA1 (N6, sc-265, Santa Cruz), b-actin (ab8227, abcam), MCM5 (Bethyl Laboratories, Inc., A300-195A-M), MCM6 (Bethyl Laboratories, Inc., A300-194A),

MCM2 (Bethyl Laboratories, Inc., A300-191A), PCNA (PC10) (Santa Cruz, sc-56), IL-17RA/IL-17R (R&D Systems, AF448).

Western blot membranes were quantified using the BIORAD Imaging system and Image Lab software.

##### Intracellular signaling by Stat3 and Stat5

Freshly harvested bone-marrow cells were enriched for Lin^-^Ter119^-^ cells using magnetic beads, as described above. The enriched cells were incubated a cytokine-free, low serum medium (IMDM with 2% FCS) for 3 hours. EPO (0.5U/ml) only, IL-17a (20ng/ml) only, or EPO (0.5U/ml) and IL-17a (20ng/ml) were then added to the medium for either 30 or 60 minutes. Cells were harvested, washed with PhosphoWash Buffer ^21^, stained with LIVE/DEAD kit (Invitrogen), fixed and permeabilized with Cytofix/Cytoperm Buffer (BD 554722) supplemented with 1mM Sodium Orthovanadate (Sigma 450243-10G), 1mM β-glycerophosphate (Sigma G9422-10G) and 1ug/ml Microcystin (EMD Millipore 475815-500UG), and Perm/Wash Buffer I (BD 557885), and frozen in freezing medium (90% FCS, 10% DMSO, 1mM Sodium Orthovanadate, 1mM β-glycerophosphate and 1ug/ml Microcystin). When thawed, cells were re-fixed and permeabilized, incubated with 5% milk and 200ug/ml Rabbit IgG (modified from Porpiglia et al., PLoS Biol. 2012), and stained with p-Stat3-AF488 (B-7) (Santa Cruz sc-8059 AF488), p-Stat5-AF647 (pY694) (BD Bioscience 612599), CD71-PE/Cy7 (Biolegend 113812), CD55-PE (Biolegend 131804), CD105-Pacific Blue (Biolegend 120412), CD150-BV650 (Biolegend 115931), CD49f (Itga6)-PE/Dazzle 594 (Biolegend 313626), CD41-BV605 (Biolegend 133921), CD117 (Kit)-APC/H7 (BD Bioscence 560185), Strepavidin-AF700 (Invitrogen S21383) and DAPI. Analysis was on an LSRII FACS analyzer.

## Extended Data Figures

**Extended Data Figure 1.**
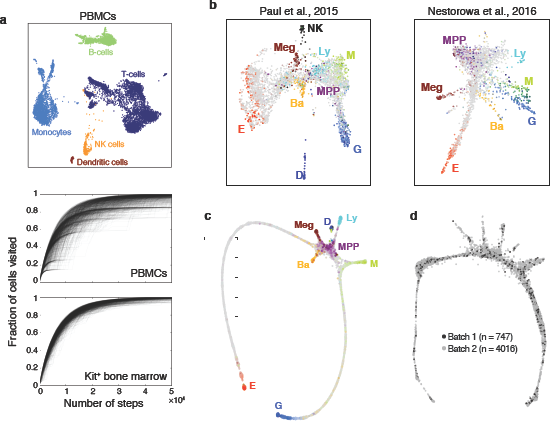
Single-cell RNA-seq of Kit+ hematopoietic progenitors for prediction of the early hematopoietic hierarchy. **a** *Upper panel:* SPRING plot of 7,959 human peripheral blood mononuclear cells (PBMCs) from 10X Genomics [https://support.10xgenomics.com/single-cell-gene-expression/datasets/2.0.1/pbmc8k]. Clusters were generated by performing spectral clustering on the underlying k-nearest-neighbor (kNN) graph and annotated on the basis of marker genes. NK cell, natural killer cell. *Middle and lower panels:* Random walks over kNN graphs for the PBMC *(middle)* and Kit+ bone marrow *(lower)* datasets. Each plot shows the fraction of nodes (cells) visited for 1,000 simulated random walks. **b** *Left panel:* SPRING plot of 2,855 lineage (Lin)^_^Kit^+^Sca1^_^ mouse hematopoietic progenitor cells from Paul et al.^22^ *Right:* SPRING plot of 1,656 cells from three mouse hematopoietic progenitor populations (Lin^-^Kit^+^Sca1^-^, Lin^-^Kit^+^Sca1+, and Lin^-^ Kit^+^Sca1+Flk2^-^CD34+) from Nestorowa et al.^23^ Colored (non-gray) cells indicate expression of lineage-specific genes (see **Extended Data Table 7**). E, erythroid; Ba, basophil/mast cell; Meg, megakaryocyte; MPP, multipotent progenitor; Ly, lymphocyte; NK, natural killer cells; M, monocyte; G, granulocyte; D, dendritic cell. **c** SPRING plot of basal bone marrow Kit+ cells (Fig. 1) constructed using only the PBA-predicted fate probabilities and differentiation ordering as inputs. Colored cells indicate expression of lineage-specific genes as in Fig. 1b. **d** SPRING plot of basal bone marrow Kit+ cells (Fig. 1), with cells colored by library preparation batch.

**Extended Data Figure 2.**
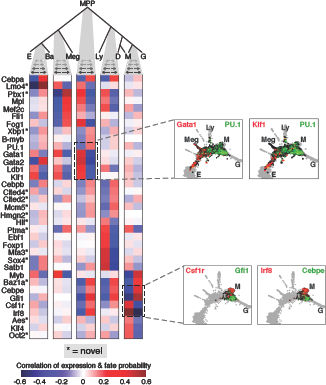
Predicting key regulators at hematopoietic choice points. Candidate regulators of fate choice, identified by ranking transcription factors and transmembrane receptors by their correlation with PBA-predicted fate probabilities at key choice points in hematopoiesis. Top-ranked genes are shown; these include many canonical regulators. Candidates not previously reported are marked by asterisk. Several candidates participate in more than one fate choice. Insets show SPRING plots colored by expression of representative gene pairs.

**Extended Data Figure 3.**
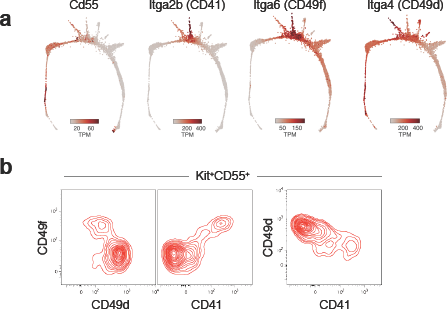
Cell surface markers used to derive a flow-cytometric sorting strategy for HPC subsets P1 to P5 (see also Fig. 2). **a** SPRING plots illustrate differentiation stage and lineage-specific expression patterns of *Cd55* and the alpha integrin chains *Itga2b* (CD41), *Itga6* (CD49f), and *Itga4* (CD49d). **b** Flow cytometric analysis of alpha integrin expression in bone marrow Kit+CD55+ cells shows a positive correlation between expression of CD49f *(Itga6)* and CD41 *(Itga2b),* and negative correlations between either of these alpha integrin chains and expression of CD49d *(Itga4).*

**Extended Data Figure 4.**
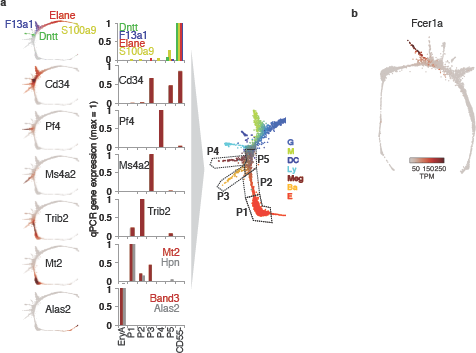
Bulk validation of mapping HPC subsets P1-P5 to the Kit+ SPRING plot. **a** Subpopulations P1 to P5 map onto specific regions of the SPRING plot. On the left are SPRING plot heat maps for a panel of marker genes; on the right are corresponding measured expression for each of the marker genes by RT-qPCR, performed on sorted cell subsets P1 to P5, and on EryA (= cells undergoing ETD^24^). A cartoon illustrates the mapping of each of P1 to P5 onto the SPRING plot based on the RT-qPCR results. RT-qPCR results are mean of at least 2 experiments, normalized to (3-actin mRNA. **b** SPRING plot heat map of expression of *Fcerla.*

**Extended Data Figure 5.**
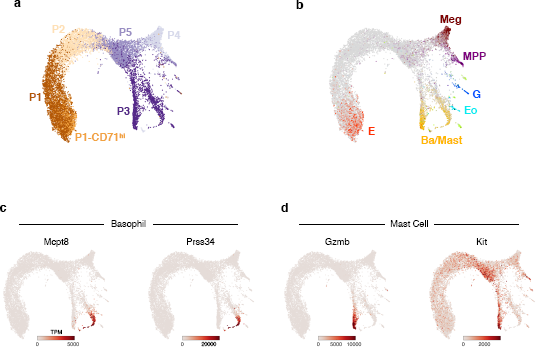
Single-cell RNA-Sequencing of HPC subsets P1-P5. **a,b** SPRING plot of single-cell transcriptomes from freshly sorted P1-P5 subsets (Fig. 2a). Cells are colored by sorted subpopulation (a) or by expression of lineage-specific marker genes (b) (**Extended Data Table 7**). E, erythroid; Ba/Mast, basophil/mast cell; Eo, eosinophil; G, granulocyte; MPP, multipotent progenitor; Meg, megakaryocyte. **c,d** SPRING plots of P1-P5 subset cells, colored by expression of basophil (c) and mast cell (d) marker genes. The larger cell number of cells in the P3 region of the graph resolves a split between the two lineages that was not observable in the original Kit+ dataset.

**Extended Data Figure 6.**
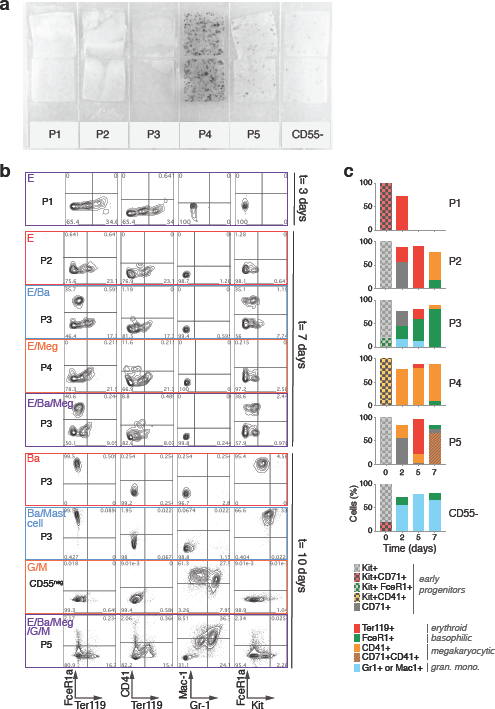
Validation of PBA predictions. **a** Megakaryocytic colonies derived from freshly sorted subsets P1 to P5 and from Kit^+^CD55^-^ cells, stained for the megakaryocytic marker acetylcholinesterase. **b** Representative flow cytometry plots to assay fate output of single cells in liquid culture (see Fig. 3). Each row corresponds to a single clone, with the left column indicating the source subset (P1 to P5, CD55^-^) of the clone and the cell type(s) produced, as inferred from the FACS plots in the remaining columns (E, erythroid, Ter119+; Ba, basophil, FcsR1a +Kit^low^; Meg, megakaryocyte, CD41+; Mast cell, Fcer1a+Kit^high^; G, granulocyte, Gr-1+; M, monocyte, Mac-1+). **c** Liquid cultures of freshly sorted P1 to P5 subsets and Kit+CD55^-^ cells in the presence of Epo and a cocktail of cytokines supporting myeloid progenitors. On the indicated days, cells were labeled with antibodies for Kit, CD71, Ter119, CD41, FcsR1a, Gr-1 or Mac-1, and analyzed by flow cytometry.

**Extended Data Figure 7.**
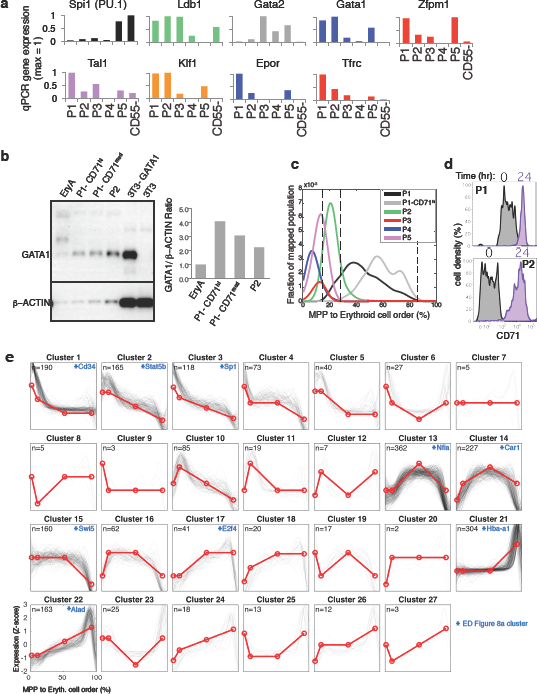
The early erythroid trajectory. **a** qRT-PCR for expression of established erythroid regulators in sorted P1 to P5 subsets. Expression of each gene is normalized to β-actin and is the mean of at least two experiments. **b** Western blot analysis for GATA1 expression in sorted P1 (subdivided into CD71^med^ and CD71^hi^ subsets), P2, and EryA (CD71+Ter119+FSC^hi^ cells, representative of ETD). **c** Density of FACS subsets P1 to P5 along the erythroid trajectory. Single-cell transcriptomes from each subset were mapped to their most similar counterparts in the Kit+ data (Fig. 2). Shown here is the fraction of mapped cells, following smoothing with a Gaussian kernel. **d** Distribution of CD71 expression in P1 and P2 cells immediately following sorting (gray) and after 24 hours of in vitro differentiation (lavender). **e** Dynamically varying genes along the MPP to erythroid axis were clustered based on their behavior across three transition points. At each transition, gene expression is either increased, decreased, or unchanged, giving a total of 27 potential dynamic patterns across all 3 transitions, shown in red. The number of genes corresponding to each dynamic pattern is noted, and individual gene Z-score normalized expression traces are shown in black. Selected clusters shown in **Extended Data Figure 8** are marked with an asterisk and a representative gene.

**Extended Data Figure 8.**
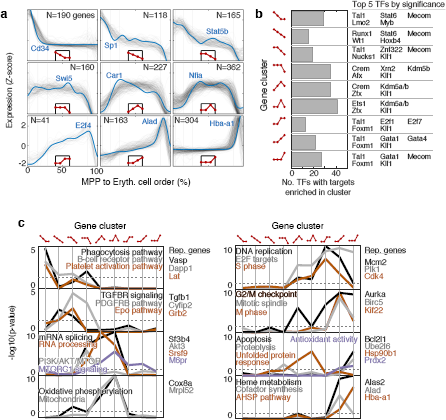
Gene set enrichment on dynamic gene cluster in early erythroid differentiation. **a** Nine key dynamic gene clusters along the MPP to erythroid progression are analyzed further for gene ontology. The clusters here are reproduced from **Extended Data Fig. 7**, with the inset in each panel indicating the dynamic pattern, used to denote the clusters in the remainder of this figure. Each node represents a progenitor stage (in order, MPP/ EBMegP+EEP/CEP/ETD), connected to the next stage by an edge that is either going up (for increased expression) or down (for decreased expression). For each cluster, a representative gene is highlighted in blue. The number of genes in each cluster is indicated. **b** Number and identity of transcription factors (TFs) whose targets are enriched in the dynamic clusters, as predicted by ChIP-X experiments^13^. **c** Significance of enrichment for signaling pathways and Gene Ontology gene sets in the dynamic gene clusters.

**Extended Data Figure 9.**
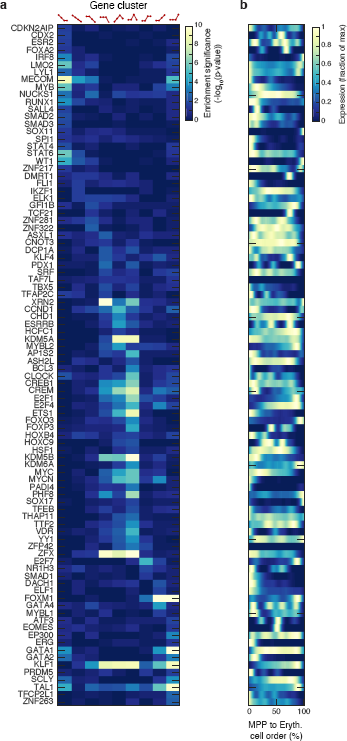
Enrichment of transcription factor targets (TFT). **a** Heatmap (-log10 of p value) of target gene enrichment for TFs (rows) with targets significantly enriched (p<0.05) in at least one of the nine dynamic clusters (columns, labeled on top) highlighted in **Extended Data Figure 8a**. Note that the TFTs shown are based on previous ChIP-X experiments^13^ and it is possible that unappreciated TFTs occur in early erythropoiesis. **b** Gene expression traces over the erythroid trajectory for the TFs from (a). Rows match those in (a).

**Extended Data Figure 10.**
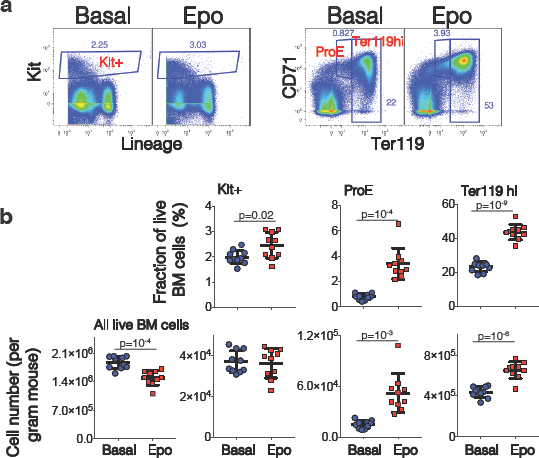
Absolute quantification of bone-marrow Kit+ cell number following ***in vivo*** administration of Epo. Adult female mice, eight weeks of age, were injected with either Epo (100 U/25 g) or saline (=basal), once per day for two days. Bone marrow was harvested at 48 hours. Viable (trypan blue negative) cells were counted using a TC20^TM^ automated cell counter (BIORAD) and stained for Kit, Ter119 and CD71 and lineage markers. **a** Flow cytometric analysis of representative bone marrow samples, gating on Kit+ Lin^-^ cells (left panels) or on proerythroblasts (ProE) and Ter119^hi^ cells (right panels; ProE and Ter119hi cells are sequential stages of ETD). **b** Data summary for two independent experiments, each with 5 mice in each group, either basal or Epo injected. Top panels show the fraction of all BM cells for each of the flow cytometric gates defined in (a). The lower panels show the absolute cell count in adult bone marrow for subsets defined as in each flow cytometric gate, or for the total number of bone-marrow cells. P values *(2-tailed t test, unequal variance)* are shown for all *p<0.05.*

**Extended Data Figure 11.**
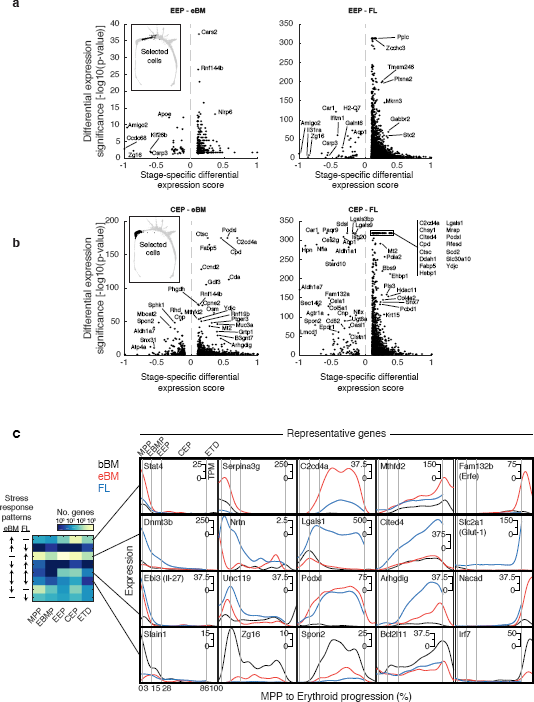
Identification of stage-specific differential gene expression during the erythroid stress response. **a** Identification of genes differentially expressed in EEP cells of either eBM (left panel) or FL (right panel), compared with basal BM. P-values were calculated using a binomial test for differential expression (see Methods) and measure the significance of the expression difference. The specific enrichment score (also described in Methods) measures the degree to which the differential expression is specific to this region of interest (EEPs); positive scores correspond to region-specific upregulation, and negative to region-specific downregulation. Selected genes are highlighted. **b** The same analysis applied to the CEP stage. **c** Stage-specific differential gene expression during stress, comparing eBM and FL. The heatmap (left) shows the number of DE genes at each stage that show similar or different patterns of upregulation and downregulation in FL and eBM. Representative gene traces are shown on the right.

**Extended Data Figure 12.**
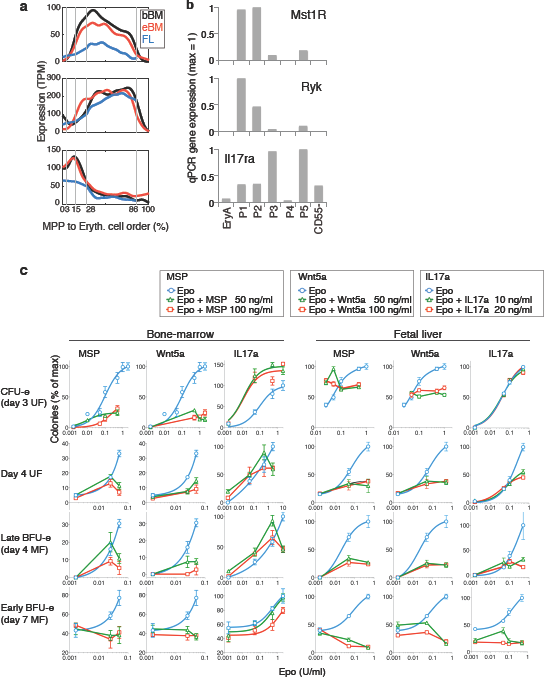
Localized gene expression and functional response of the erythroid lineage to stimulation of Mst1, Ryk and IL-17Ra. **a,b** Predicted expression pattern (a) and confirmation by RT-qPCR (b) for *Mst1r, Ryk* and *Il17ra* in basal BM. In (a) traces show the smoothed scRNA-Seq gene expression on cells arranged along the erythroid trajectory in basal BM, FL and eBM. **c** Complete results for CFU-e colony formation assays in methylcellulose, supporting the data shown in Fig. 5. Curves show CFU-e numbers in the presence of increasing concentrations of Epo, or Epo with added ligand, either MSP, Wnt5a or IL-17a. Error bars show SD of two independent experiments, with four replicates per experiment. Where appropriate, data was fitted with a dose-response curve, Hill coefficients.

**Extended Data Figure 13.**
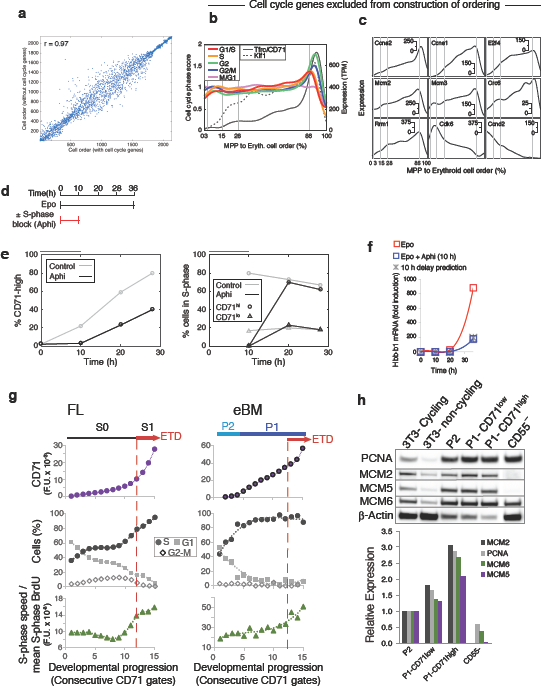
Independence of cell ordering on cell cycle genes, and evidence of an S-phase dependent CEP-to-ETD transition in BM erythropoiesis. **a** The computational ordering of cells from MPP to ETD is not sensitive to whether or not annotated cell cycle genes are included (cell ordering correlation is R=0.97). **b,c** Reproduction of main text figure panels 6b,c after excluding cell cycle genes shows that the computationally inferred gene expression dynamics of cell cycle genes during EEP/CEP differentiation are not sensitive to whether or not annotated cell cycle genes are included in ordering cells. **d-f** Activation of ETD is dependent on S phase. **d** Schematic illustration of experiments testing the link between S-phase progression and the CEP-to-ETD transition. BM Kit+Lin^-^CD71^-^ cells were cultured in the presence of Epo for 36 hours, and either in the presence or absence of the DNA polymerase inhibitor Aphidicolin (Aphi) for the first 10 hours. **e** BM Kit+Lin^-^CD71^-^ cells require S phase in order to upregulate CD71, an initial event in ETD. Cells were treated as in (d); *left:* CD71^hi^ cells fail to appear in the first 10 hours if cells are exposed to Aphi; they appear as soon as Aphi is removed from the medium. *Right:* cell cycle analysis of the same cells shows that Aphi prevented S phase progression during its presence in the culture medium; Aphi removal was followed by full recovery of S phase progression, with a high fraction of CD71^hi^ cells in S phase. **f** Aphi exposure for 10 hours delays induction of b-globin *(Hbb-b1)* by 10 hours. **g** CD71 expression (top row), cell cycle phase distribution (middle), and intra-S phase DNA synthesis rate (lower), for consecutive FACS gates of increasing CD71 in early stages of erythropoiesis from the fetal liver (left panel) and Epo-simulated bone marrow (right). See **Fig. 6e,f** for similar analysis in bBM. **h** Western blots (upper panel) and their quantification by densitometry (lower) showing an increase in S phase proteins during progression from EEP/P2 to early CEP (P1-CD71^b^ to late CEP (P1-CD71^hi^). Controls 3T3 cells were either cycling, or contact-inhibited (non-cycling), as indicated.

## Extended Data Table Legends

### Extended Data Table 1 Lineage-specific marker genes

List of marker genes for mature cells of each lineage and multipotent progenitors. These genes were used for visualizing cell types on SPRING plots and for identifying the most mature cells for Population Balance Analysis. Marker gene lists are sample-specific due to sample-to-sample differences in the detected lineages and the maturity of these lineages.

### Extended Data Table 2 Genes correlating with branching computationally predicted fate probabilities in hematopoietic progenitor cells

The table shows the process by which genes were selected, as well as their identity. Column 1 (Cell subset) shows the PBA gates used, and for each cell subset, Column 2 (Lineage bias score) indicates the score to which the genes identified were correlated. Columns 3-5 give the Gene symbol, its Correlation to the lineage bias score in the cell subset, and the P-value of the correlation. For example, “P(Er)*P(Ba) > 0.07” identifies cells computationally predicted to reside close to the Erythroid-Basophil branch point, as measured by the product of the PBA scores P(Er) and P(Ba). Within this subset genes were tested for correlation against the PBA score “P(Er)-P(Ba)”. Thus genes with positive correlation to this score were Erythroid biased and genes with negative correlation were Basophil biased.

### Extended Data Table 3 Dynamic changes in gene expression along the erythroid trajectory

List of significantly varying genes in the basal BM dataset. For each gene, maximal and minimal expression levels, and the corresponding locations along the MPP to erythroid progression, are shown. See methods section for calculation of FDR-corrected p value.

### Extended Data Table 4 Gene set enrichment analysis (GSEA) of dynamic gene clusters

Genes assigned to each dynamic cluster (sheet 1), and GSEA results for signaling pathways and Gene Onotology terms (sheet 2) and transcription factor targets (sheet 3). Dynamic gene clusters are numbered as in **Extended Data Fig. 8**. In the GSEA tables (sheets 2 and 3), enriched gene set titles (column B), corresponding corrected p value (column C) and gene hits (column D) are shown.

### Extended Data Table 5 Genes differentially expressed in stress

Genes differentially expressed (DE) in each of the MPP, EBMP, EEP, CEP, and ETD stages are shown, in two separate sheets for eBM and FL, respectively. See methods section for calculation of the DE score (column C).

### Extended Data Table 6 Genes correlated with progression through the CEP stage

For each gene expressed in the CEP stage, this table lists the “Linearity score”, a measure designed to identify genes that steadily increase or decrease with progression along the differentiation ordering (see Methods section “Identifying genes that change steadily in the CEP stage”). Large positive scores indicate genes that are strongly up-regulated in a linear fashion throughout the CEP stage; large negative scores indicate genes that are down-regulated linearly; and scores close to 0 indicate genes whose expression changes little or in a non-linear fashion.

### Extended Data Table 7 Modifications to the inDrops protocol

Detailed list of changes to the inDrops protocol by sample.

